# Industrialization drives convergent microbial and physiological shifts in the human metaorganism

**DOI:** 10.1101/2025.10.20.683358

**Authors:** Mathilde Poyet, Malte Rühlemann, Ana P. Schaan, Yue Ma, Lucas Moitinho-Silva, Eike M. Wacker, Hannah Jebens, Lucas Patel, Le Thanh Tu Nguyen, Alexis Zimmer, Damian Plichta, Daniel McDonald, Christine Stevens, Adwoa Agyei, Mary Y. Afihene, Shadrack O. Asibey, Yaw A. Awuku, Aida S Badiane, Lee S. Ching, Chris Corzett, Awa Deme, Manuel Dominguez-Rodrigo, Amoako Duah, Alain Fezeu, Alain Froment, Sean Gibbons, Catherine Girard, Jeff Hooker, Fatimah Ibrahim, Deborah Iqaluk, Vanessa Juimo, Pinja Kettunen, Sophie Lafosse, Ernest Lango-Yaya, Jenni Lehtimäki, Yvonne A. L. Lim, Audax Mabulla, Varocha Mahachai, Rihlat S. Mohamed, Katya Moniz, Ivan E. Mwikarago, Yvonne A. Nartey, Daouda Ndiaye, Mary Noel, Charles Onyekwere, Tan M. Pin, Amelie Plymoth, Lewis Roberts, Lasse Ruokolainen, John Rusine, Laure Segurel, B. Jesse Shapiro, Shani Sigwazi, Ainara Sistiaga, Kenneth Valles, Tommi Vatanen, Ratha-korn Vilaichone, Philip Rosenstiel, John Baines, Andre Franke, David Ellinghaus, Rob Knight, Mark Daly, Ramnik J. Xavier, Eric J. Alm, Mathieu Groussin

**Author notes:** Corresponding authors: Mathieu Groussin, Mathilde Poyet and Eric Alm. co-first authors (Mathilde Poyet, Malte Rühlemann). co-second authors (Ana P. Schaan, Yue Ma, Lucas Moitinho-Silva, Eike M. Wacker). These authors contributed equally to this work. co-senior authors (Eric J. Alm, Mathieu Groussin).

## Abstract

Understanding how host lifestyle and industrialization shape the human gut microbiome and intestinal physiology requires multimodal analyses across diverse global host contexts. Here, we generate multivariate data from the Global Microbiome Conservancy cohort, including gut microbiome, IgA-sequencing, host genotyping, diet, lifestyle and fecal biomarker profiles, to investigate host–microbiome interactions across gradients of industrialization and geography. We show that industrialization is associated with homogenized microbial compositions, reduced microbial diversity, and lower community stability, independent of host confounders. We further show that industrialization is linked to elevated markers of gut stress, increased IgA secretion, and altered patterns of IgA-bacteria interactions. Finally, we show that microbiome-based disease predictors trained on industrialized populations lose accuracy in less industrialized cohorts, highlighting limited cross-population transferability. Together, our results suggest profound restructuring of host-microbiome interactions due to industrialized lifestyles, and emphasize the need for inclusive, globally representative data to improve translational microbiome applications across diverse human populations.

## Introduction

The transition to industrialized lifestyles has been linked to marked changes in the human gut microbiome, including reduced microbial diversity, pronounced shifts in microbial compositions, and distinct metabolic and virulence capacities ^1–21^. The extent and potential mechanisms by which host factors, such as host diet, genetics, geography, immune response and local infectious burden, confound these patterns remain unclear ^22,23^. In addition, to what extent these patterns result from inconsistent methods for sample collection, preservation and data generation across populations and studies is unknown ^24^. Furthermore, whether industrialization-associated differences in microbial diversity and composition impact ecological characteristics of the microbiome, such as community stability, or impact host immune and inflammatory responses, is unknown. As such, understanding global human gut microbiome variation and its impact on host-microbiome systems requires the recruitment of multiple and diverse populations from worldwide regions along gradients of industrialization and geographic distances. It also necessitates the collection of multi-dimensional host data and using consistent protocols for sample collection and data generation.

To address these limitations, we present the Global Microbiome Conservancy (GMbC) cohort and data. We provide multiple lines of evidence that lifestyle, and industrialization in particular, shapes the human metaorganism, i.e. the integrated unit of the human host and its associated microbiota. These effects occur independently of host confounders and manifest as shifts in microbiome diversity, composition, and stability, as well as in host immune responses and homeostasis.

In studying the microbiomes of human populations with diverse lifestyles worldwide, we have found that common terminology in the literature, including our own publications ^25^, can be unsatisfactory ^26^. For instance, the terms used to describe subsistence strategies or lifestyles often fall short of accurately capturing the complexities of the human populations that we collaborated with in the field. Additionally, some of these terms carry assumptions that may reflect biases rooted in colonial histories or imply that these populations have been static during human history and evolution ^27^. As such, they risk perpetuating misconceptions and stereotypes ^28^. For example, labels like “hunter-gatherer” fail to capture the diversity of subsistence practices within these populations and overlook their recent cultural history ^29^. We recognize that no terminology is entirely neutral or capable of fully representing the complexities of human societies. Key terms, including industrialization, urbanisation and subsistence strategies are thus defined, with associated limitations, and potential biases in the Methods and Supplementary Information (Table 1). With this approach, we aim not only to support accurate interpretation of our data, but also to encourage critical reflection and to contribute to a less biased, inclusive framework for understanding changes in global microbiome diversity and function in the context of industrialization.

## Results

### Global and multi-factor profiling of the human metaorganism across geographic scales

We hypothesize that global and multi-factor profiling of human metaorganisms will allow us to begin teasing apart the relative contributions of host factors – such as industrialization, diet, and genetics – to microbiome composition and function. The GMbC cohort consists of adult participants (n = 1,015) from 12 countries, with a mean age of 36.7 years (range: 18–87), a mean BMI of 22.8 (range: 12.9–44.7), and a balanced sex ratio (51.2% female, 48.9% male) (Fig. 1A-C). All participants were recruited without self-reported signs of infection or chronic diseases. Our recruitment strategy was designed to enable comparisons across multiple geographic scales ranging from regional to local while accounting for differences in lifestyle (including industrialization, urbanization and subsistence strategies), diet and host genetics (Supplementary Table 1). To capture fine-scale host-microbiome variation, we recruited multiple populations within 11 of the 12 countries, spanning 35 localities in total (Fig. 1B & Supp. Table 1). This approach resulted in pairwise geographic distances between participants ranging from 0 to 14,920 km (Fig. 1D). Participants reside in areas with population densities ranging from 0.4 to 15,000 inhabitants per km² (Supp. Table 1).

**Figure 1.**
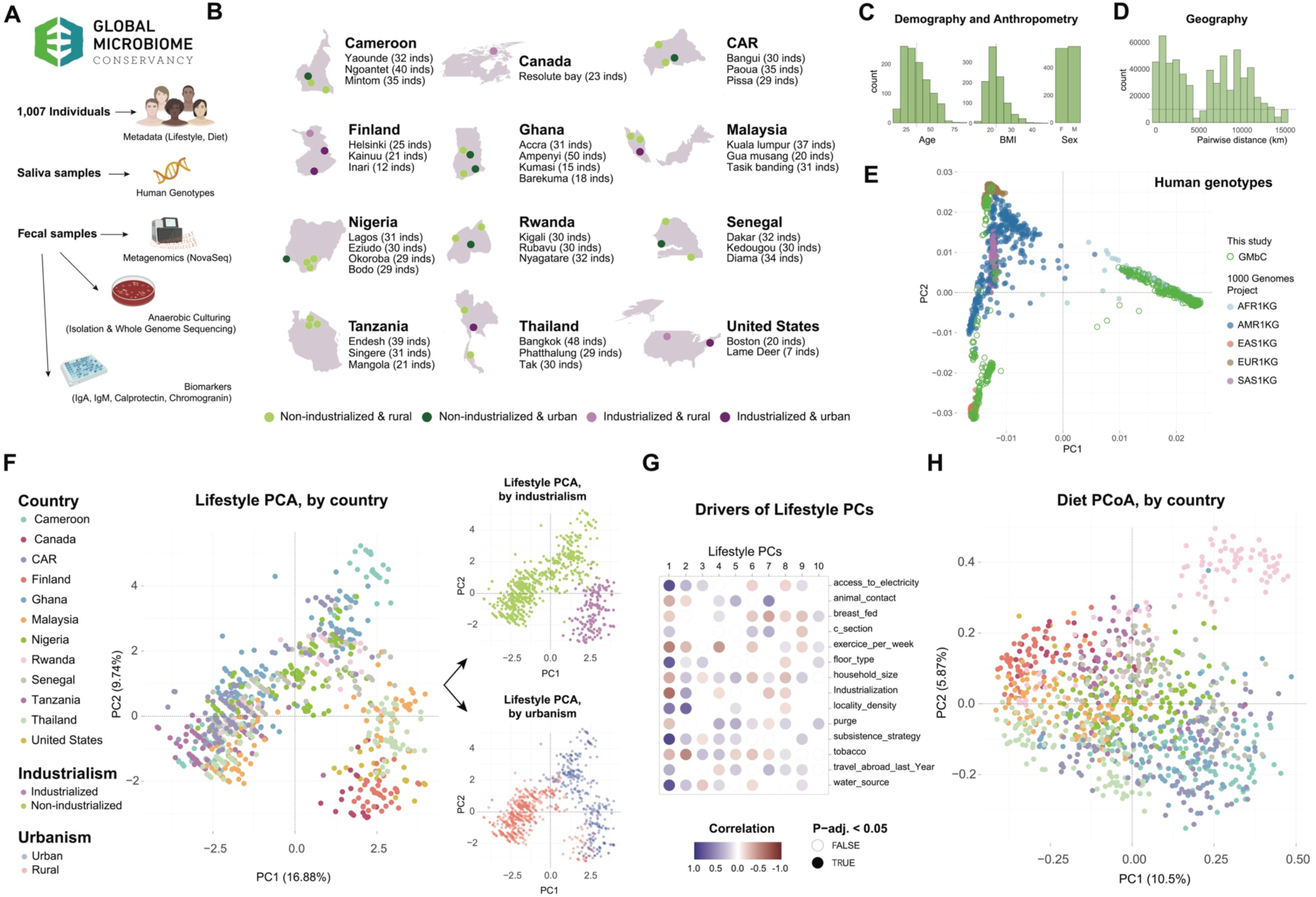
Multimodal sampling in the GMbC cohort to investigate factors shaping human metaorganisms worldwide. A. Collected samples and metadata from GMbC participants (n = 1,015). Gut shotgun metagenome, human genotype, and cultured isolate genomic short read data were generated. Levels of fecal biomarkers (IgA, IgM, calprotectin and chromogranin) were measured. Lifestyle and diet metadata were collected. B. Countries (n = 12) and sampling locations (n = 35). Locations are colored based on industrialism and urbanism status of recruited populations. The number of participants per location is shown. C. Distribution of age, BMI and sex parameters in the GMbC cohort. D. Distribution of pairwise geographic distances across sampled participants (Haversine distance, assuming a spherical Earth). E. PCA of human genotypes of the GMbC cohort (green circles) and the 1000 Human genomes project (plain points, colors represent superpopulations). F. Left plot: Factor analysis of mixed data (FAMD) applied to lifestyle metadata factors (see Methods). Participants are colored by country. Top right: same plot, participants colored by industrialism status. Bottom right: same plot, participants colored by urbanism status. G. Correlation between individual lifestyle factors and the first ten principal components (PCs) of the Lifestyle PCA. Statistically significant associations (adjusted p-val < 0.05) are shown in plain circles. H. Principal component analysis (PCoA) of diet metadata (UniFrac-based distances of FFQ data, see Methods). Participants are colored by country as in F, left plot.

We generated shotgun metagenomic sequencing data (Fig. 1A) to capture the taxonomic and functional composition of the gut microbiome (average of 25.5M reads per sample; average mapping rate of 85% (sd = 3.7%); see Methods, Supp. Fig. 1A and Supp. Table 1). The cohort is composed of individuals from 73 self-declared ethnicities. We generated human genotyping data (n = 913) to contextualize microbiome variation with host genetic diversity, and confirmed that GMbC participants are from diverse genetic backgrounds (comparison with data from the 1000 Human Genome Project, Fig. 1E). We further assessed global population structures, and assigned individuals to 17 admixture groups (see Methods and Supp Fig. 1B).

We collected comprehensive lifestyle and dietary metadata (see Supp. Table 1), and applied dimension reduction approaches (see Methods) to extract the main metadata variation components (PCs) for use in variance partitioning of microbiome data (see below) (Fig. 1F & H). For clarity, we use the capitalized terms *Lifestyle*, *Genetics*, and *Diet* to refer to the main PCs derived from these respective metadata sets. Several factors contribute to the two first PCs of lifestyle, including industrialization status, access to electricity, subsistence strategy and household floor type for PC1_Lifestyle, and population density and tobacco usage for PC2_Lifestyle (Fig. 1F & G). **Importantly, PC1_Lifestyle captures the main variation associated with societal industrialization (Fig. 1F), with higher PC1_Lifestyle values reflecting more industrialized societies. We use PC1_Lifestyle throughout the paper as a continuous proxy to assess the impact of industrialized lifestyles on the microbiome.** The main items contributing to dietary variations include dairy products such as milk, cheese and yogurt, and industrialized goods such as ice cream for PC1_Diet, and macabo, guava and wild mammals for PC2_Diet (Supp. Fig. 1C).

As expected, global lifestyle variation is partly correlated with geography, Genetics, and Diet. For example, PC1_Lifestyle shows collinearity with geographic variables (latitude and longitude), genetic factors (PC1 and PC2), and, to a lesser extent, with diet (PC1) (Supp. Table 1).

Overall, the GMbC dataset represents a unique resource of integrated microbiome, lifestyle, dietary, genetic, and environmental data across globally sampled cohorts. In the next two sections, we leverage this resource to characterize microbiome variation, disentangle the contribution of host factors, and interrogate the effect of industrialization on the microbiome.

### Host lifestyle and industrialization, independently of host confounders, drives loss of microbiome diversity and homogenization of microbial compositions

We first aimed at quantifying associations between microbial diversity and host factors. We found that alpha taxonomic diversity is negatively correlated with PC1_Lifestyle (Spearman test, rho = -0.44, p-val = 2.6e-47), with individuals from industrialized countries and urban areas hosting lower microbial diversity in their microbiome (Fig. 2A) (alpha diversity measured with Faith’s phylogenetic diversity index on species-level genome bin (SGB) profiles; similar trends observed with Shannon index (Supp Table 2)). This trend is also replicated at higher geographic resolution, within individual countries (Supp. Fig. 2A and Supp Table 2), suggesting convergent decreases of microbiome diversities worldwide. We further found that alpha diversity is correlated to host Diet (PC1_Diet, rho = -0.38, p-val = 4.9e-34), Genetics (PC1_Genetics, rho = 0.17, p-val = 9.9e-07), geography (latitude, rho = -0.33, p-val = 1.6e-25), age (rho = 0.17, p-val = 2.0e-07), BMI (rho = -0.15, p-val = 2.0e-06), and sex (Wilcoxon test, 53.8 (mean Female) vs. 48.4 (mean Male), p-val = 4.5e-05).

**Figure 2.**
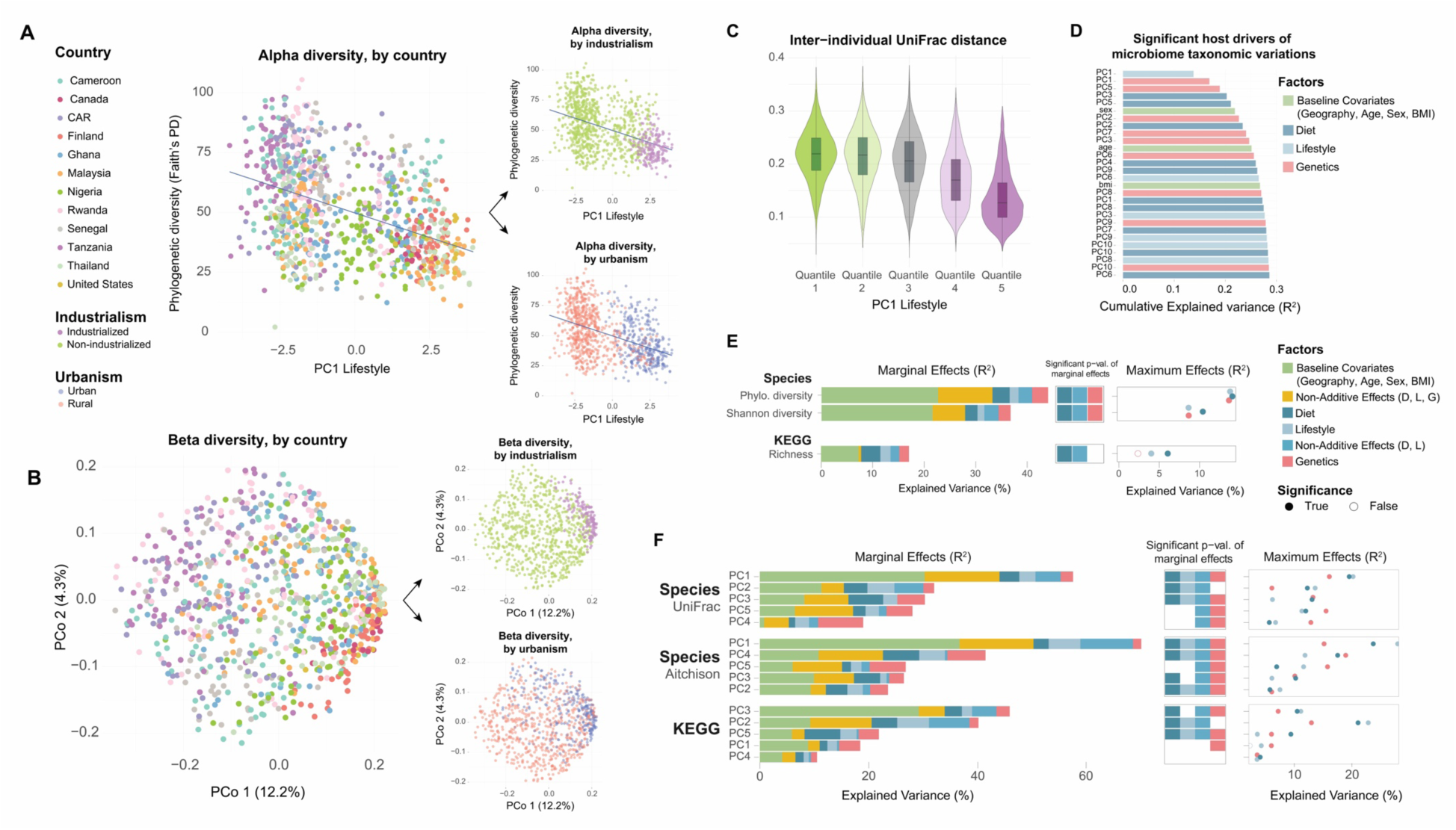
Effect of industrialized and urban lifestyles on community-level variations in the global human gut microbiome. A. Phylogenetic diversity (Faith’s PD index) of metagenomes (species level) as a function of the first PC of the Lifestyle PCA. PC1 Lifestyle is used as a continuous proxy for industrialization (Fig. 1F&G). Participants are colored by country (left plot), industrialization status (top right) or urbanism status (bottom right). The blue line shows the linear regression. B. PCoA of unweighted UniFrac compositional dissimilarities (species level). Participants are colored by country, industrialism status and urbanism status as in A. C. Inter-individual compositional dissimilarities across quantiles of PC1 Lifestyle, used as a continuous proxy for industrialization (Fig. 1F&G), increasing from first to fifth quantile. Industrialized and urban populations exhibit increased homogeneity of bacterial compositions (quantiles #4 & #5). D. Cumulative explained variance (as measured by stepwise dbRDA, forward multivariate model selection) of species-level microbiome compositions by individual host factors. The baseline covariates (latitude, longitude, age, sex and BMI) and the first 10 PCs for Lifestyle, Diet and Genetics were included in the model. E. Variance partitioning of alpha diversity of taxa (species level) (Faith’s PD and Shannon diversity) and KEGG KO functions (Richness). Marginal and maximum variances (R^2^) of alpha diversity were calculated for the following factors: Lifestyle (10 first PCs), Diet (10 first PCs), Genetics (10 first PCs), Non-Additive Effects of Lifestyle and Diet, and Non-Additive Effects of Lifestyle, Diet and Genetics. Statistical significance is represented as plain square and circle symbols. Marginal and maximum effects are calculated using a nested model approach where the null model contains baseline host covariates (age, sex, BMI and geography), and the full model contains all tested host variables (see Methods). F. Variance partitioning of PCs of beta diversity for taxa (species level) (unweighted UniFrac and Aitchison distance) and KEGG KO functions (Canberra distance). The same approach as in panel E was used. Colors and symbols for statistical significance are as in panel E.

We next explored how host variation shapes microbiota composition. We found that compositions are associated with host lifestyle and industrialization (Fig. 2B, PERMANOVA test, SGB-level unweighted UniFrac beta diversities ∼ PC1_Lifestyle, R2 = 0.14, p-val < 0.001; similar trends are observed with Aitchison index (Supp Table 2)). PC1 of microbiome compositions separates individuals from non-industrialized societies from those in all industrialized societies worldwide, which cluster together (Fig. 2B). Microbiome composition of populations living in urban, densely populated areas of non-industrialized countries, such as Accra, Lagos, or Bangui clusters in between, in an intermediary state (Fig. 2B & Supp. Fig. 2B). In addition, inter-individual variability in compositions is negatively associated with PC1_Lifestyle (Fig. 2C, Kruskal-Wallis test, Chi-sq = 19597, p-val < 2.2e-308), showing a reduction in inter-individual variability in the microbiome of urban and industrialized societies. Overall, these findings suggest that as populations become more urbanized and societies industrialize, similar processes drive convergent reductions in microbial diversity and shift inter-individual differences toward greater homogeneity.

We also measured strong associations between compositional dissimilarities and host diet (unweighted UniFrac ∼ PC1_Diet, R2 = 0.10, p-val < 0.001), as well as with host genetics (PC1_Genetics, R2 = 0.06, p-val < 0.001). We further found a significant association with host geography, when considering latitude (R2 = 0.07, p < 0.001) and the physical distance between individuals (Haversine distance, Spearman correlation, ρ = -0.04, p = 1.8e-138; Supp. Fig. 2C). Significant associations were also measured with host age (PERMANOVA, R2 = 0.01, p-val < 0.001), BMI (R2 = 0.02, p-val < 0.001), and sex (R2 = 0.009, p-val < 0.001). Finally, we found similar trends of association between these host traits and alpha/beta diversity of both KEGG KOs and 50% similarity homologous gene families (Supp. Fig. 2D-E & Supp. Table 2; e.g. associations with PC1 Lifestyle: PERMANOVA tests; R2 = 0.078, p-val < 0.001 for KEGG categories; R2 = 0.08, p-val < 0.001 for gene families). Overall, effect sizes for Lifestyle, Diet, Genetics and Geography in the GMbC cohort are higher than the largest effect sizes typically observed for host metadata variables in population cohorts with more homogeneous lifestyle and diet ^30,31^, emphasizing the extent of variation that exists in the gut microbiome at global scales.

Next, we determined the nonredundant cumulative contribution of host factors to the unweighted UniFrac-based compositional variation. For this, we performed a forward multivariate model selection (stepwise dbRDA) with the 10 first PCs of Lifestyle, Diet, and Genetics, as well as age, sex, BMI and geography (latitude and longitude). We found that 28 of these factors were retained as having significant contributions (cut-off p-vals < 0.05), including multiple PCs from lifestyle, diet and genetics categories (Fig. 2D). Latitude and longitude were not conserved, likely due to their collinearity with PC1_Lifestyle (Supp. Table 1). The cumulative (nonredundant) effect size of these variables on microbiome compositions amounts to 28.6% (Fig. 2D).

To disentangle contributions of individual host variables on microbiome variations, we applied a variance partitioning approach on alpha and beta diversity profiles. Importantly, we measured contributions of Lifestyle, Diet and Genetics after correcting for baseline covariates, which consists of geography, age, sex and BMI. This approach is conservative, as geography (latitude and longitude) is correlated with PC1_Lifestyle and PC1_Genetics (Supp. Table. 1). We defined a “null” model where microbiome variation is solely explained by these baseline covariates, and a “full” model, which includes Lifestyle (first 10 PCs), Diet (first 10 PCs), and Genetics (first 10 PCs) in addition to the baseline covariates. We then compared models that are nested with the null model to quantify the <u>maximal</u> variance of these host factors, and models that are nested within the full model to quantify <u>marginal</u> variances, i.e. the additional fraction of variance uniquely explained by a given factor after accounting for all other factors (see Methods for a detailed description of compared models). <u>Importantly, our use of the term *marginal* throughout the</u> <u>manuscript refers to these unique contributions and should not be interpreted as implying weak</u> <u>or negligible effects.</u> We also quantified the marginal effects of lifestyle and diet combined (hereafter referred to as the ‘Environment’ effect), and of the entangled collinearity of Lifestyle, Diet, and Genetics. We found that our full model explains 44% and 57.8% of the variation (R^2^) in alpha and beta diversity, respectively (Faith’s Phylogenetic Diversity, Fig. 2E; unweighted UniFrac PC1, Fig. 2F), showing that our diverse set of host modalities captures a large fraction of microbiome variations.

We further found that Environment, Diet and Genetics have significant marginal effects to variation in alpha diversity (Fig. 2E and Supp. Table 2; Faith’s Phylogenetic diversity: R2 = 0.77, p-val = 3.32e-14; R2 = 0.033, p-val = 4.77e-7; R2 = 0.031, p-val = 2.12e-6; respectively) and beta diversity (Fig. 2F; unweighted UniFrac PC1: R2 = 0.113, p-val = 1.17e-31; R2 = 0.037, p-val = 5.15e-11; R2 = 0.023, p-val = 2.91e-6; respectively), even when accounting for all other covariates in the full model. The marginal effect of lifestyle alone is also found to be significant for compositions (Fig. 2F and Supp. Table 2, unweighted UniFrac PC1: R2 = 0.300, p-val = 1.1e-8), but not for alpha diversity, even though its maximum possible effect is in the same range as diet and genetics (Fig. 2E, Faith’s Phylogenetic diversity: R2 = 0.112, p-val = 5.22e-17). Similar trends are observed for the partitioning of variation in community-level functional diversity (e.g. KEGG KOs) (Fig. 2E & F, and Supp. Table 2). Overall, our results show that lifestyle, diet, and genetics each contribute independently and significantly to gut microbiome variation, despite their partial collinearity, as revealed by our nested multivariate modeling approach that accounts for geography.

### Disentangled effects of industrialization reveal shifts in individual microbial features and cobalamin biosynthesis enrichment

Next, we investigated the effect of host lifestyle and industrialization on individual taxa and functions. We first found that individual bacterial taxa respond consistently to specific variation in host lifestyle across human populations and geographic regions (Fig. 3A & Supp. Fig. 3A). Comparing urban and rural groups within countries, we found that taxa showed consistent differential abundance patterns across countries (Inverse-variance weighted fixed-effects meta-analysis per taxon, 71% of species with significant q-values; Fisher’s method for combining p-values: p-val < 2.2e-16; Supp. Table 3). This finding suggests the existence of common, host lifestyle-related ecological mechanisms that convergently drive shifts of individual bacteria worldwide.

**Figure 3.**
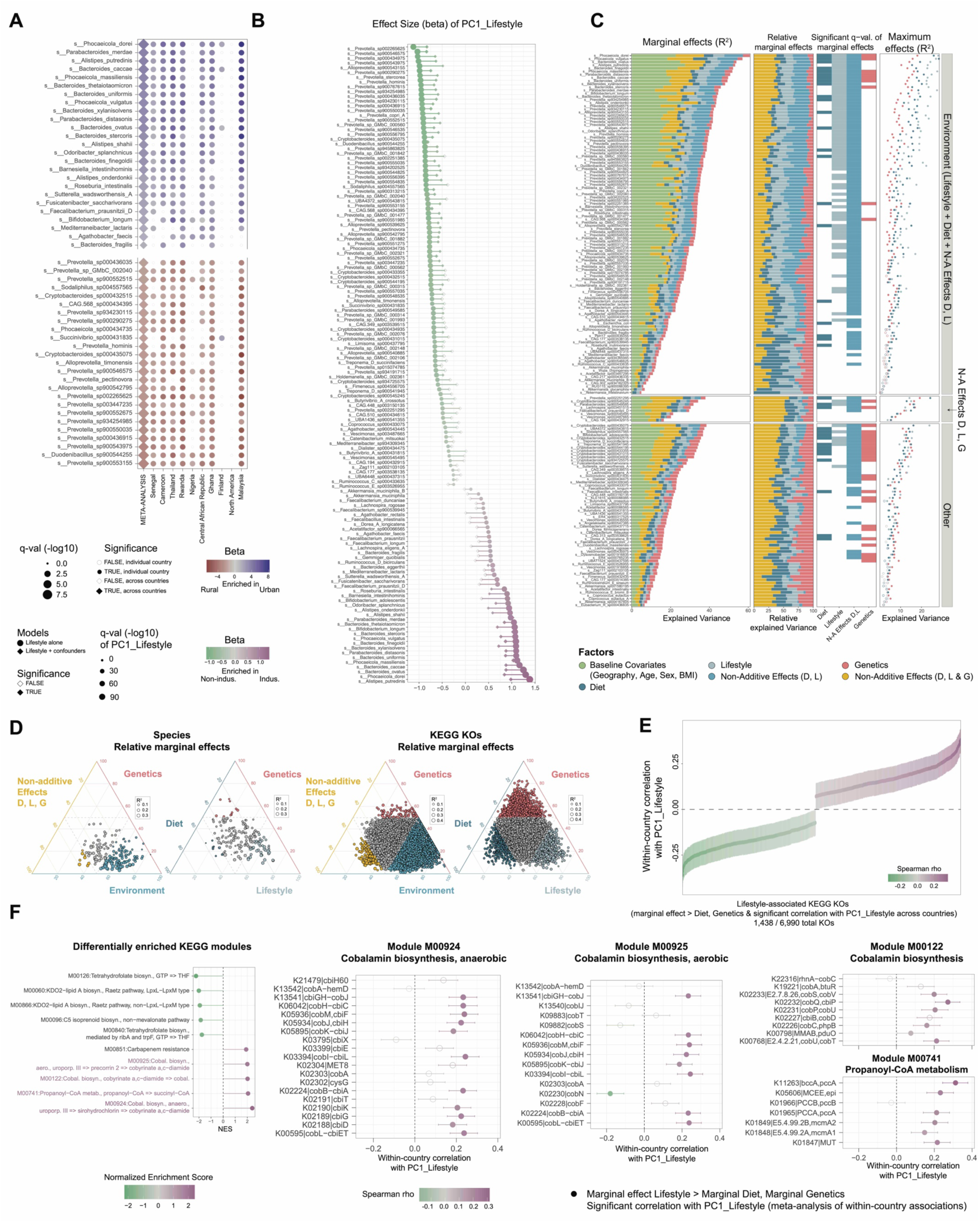
Impact of industrialized and urban lifestyles on individual taxonomic and functional features of the gut microbiome. A. Meta-analysis of country-level associations between gut bacterial species abundance and urbanism status (urban vs. rural). The “META-ANALYSIS” column shows statistical results of the cross-country meta-analysis performed using an inverse-variance weighted fixed-effects approach. Because Canada included only rural participants in our cohort, it was grouped with the USA under the label ‘North America’. Species are shown in rows. The top 25 species most enriched in each category are shown. B. Effect of host covariates on the association between bacterial species abundance and industrialization (PC1 Lifestyle) (x axis: species; y axis: effect size). Two models were compared – a first model with Lifestyle PCs alone as predictors, and a second model with host covariates (baseline covariates and Diet and Genetic PCs). Significant associations are shown as plain symbols. C. Variance partitioning of species abundance to calculate marginal and maximum variance of baseline, Lifestyle, Diet and Genetic factors, along with Non-additive Effects. The same statistical framework and figure design as in Fig. 2E&F were used. “Environment” is defined as the overall contribution of Lifestyle and Diet (see Methods). Features are grouped by majority factors that explain >50% of variance. Data for KEGG KOs is shown in Supp. Fig. 3C and Supp. Table 3. D. Ternary plots of relative marginal effects on species (left) and KEGG KOs (right) profiles. Marginal effects for KEGG KOs were calculated as for species shown in panel C (see Supp. Fig. 3). Relative contributions of Genetics vs. Environment vs. Non-additive effects of Diet, Lifestyle and Genetics are shown in left plots, while contributions of Genetics vs. Diet vs. Lifestyle effects are shown in right plots. E. Identification of lifestyle-associated KEGG KOs across countries. Lifestyle-associated KEGG KOs were identified using the following criteria: (i) significant association with either Lifestyle alone or Environment, based on marginal variances calculated as in panel C, (ii) effect size (marginal R2) for Lifestyle higher than for Genetics and Diet, (iii) significant association with industrialization status (indus vs. non-indus, q-value < 0.05), (iv) significant association (q-value < 0.05) with PC1 Lifestyle and (v) significant within-country meta-association with PC1 Lifestyle. F. KEGG module enrichment analysis reveals Cobalamin biosynthesis as enriched in the industrialized group (left panel). The analysis is based on the average country correlation of KEGG KOs with PC1 Lifestyle. KEGG modules enriched in the industrialized group and highlighted in purple are described in further panels. Panels for modules M00924, M00925, M00122 and M00741 show the distribution of within-country correlation coefficients between all individual KEGGs of these modules and PC1 Lifestyle. KEGGs with plain symbols are part of the set of Industrialization-associated KEGGs identified in panel E.

We then tested whether these lifestyle-related effects on individual taxa are influenced by confounding host factors, which was rarely addressed in prior studies. Previous work identified marker taxa of industrialization, including the “VANISH” taxa (e.g. Prevotellaceae, Succinivibrionaceae, and Spirochaetaceae), which are depleted in industrialized societies, and the “BLOSSUM” taxa (e.g. Bacteroidaceae and Verrucomicrobia), which are enriched in industrialized societies ^32^. Accounting for the influence of Diet, Genetics, Geography and other baseline covariates, we found that multiple marker taxa, such as Bacteroides, Prevotella, and Phocaeicola species, remain significantly associated with industrialization (PC1_Lifestyle) (Fig. 3B & Supp. Fig. 3B, see Methods). However, several previously identified VANISH and BLOSSUM taxa were no longer significantly associated with PC1 Lifestyle after accounting for these confounders (Fig. 3B & Supp. Fig. 3B), such as Treponema (Spirochaetes), Succinivibrionaceae, and Akkermansia (Verrucomicrobia). This suggests that the abundance of these bacteria is not primarily or solely driven by industrialization. Instead, more complex mechanisms involving other host factors, possibly in interaction with lifestyle and industrialization, contribute to the observed worldwide variation in these taxa.

We next quantified the variance in individual microbial features explained by lifestyle, diet and genetics after correcting for baseline (demographic and geographic) confounders. For this, we used the same variance partitioning approach as described above (See Methods) on taxa and functional categories that have a prevalence > 20% (Fig. 3C&D, Supp. Fig. 3C and Supp. Table 3). We found that host factors explain variation in microbial profiles in the range of 3-60% for species, and 3-64% for KEGG KOs. We further found that environment and lifestyle have a profound impact on individual features, with significant marginal effects on 74% and 40% of taxa, and 33% and 5% of KEGG KOs abundances, respectively (Fig. 3C&D and Supp. Table 3). The marginal effect (R^2^) explained by lifestyle alone exceeds that of diet and genetics for 49% of taxa and 23% of KEGG KOs (Fig. 3D). High maximum effects for lifestyle (>15% of variance) are also observed for a diverse set of Prevotella, Bacteroides and Phocaeicola SGBs (Fig. 3C). Overall, host genetics has a smaller contribution on taxa and functions (genetics has a significant marginal effect on 20% of species and 12% of KEGG KOs, Fig. 3D), confirming previous work ^33^. Notable exceptions with marginal abundance variation for Genetics that is higher than for Lifestyle and Diet are Dialister sp000434475 and Catenibacterium sp000437715 (Fig. 3C, Supp. Table 3; 6.1% and 4.0% marginal variation, respectively). These two taxa were previously found to co-diverge among Great Apes and within humans ^34^, and to be linked to specific human genetic variants ^35^.

Having partitioned the contribution of host factors on individual functional categories, we next characterized functional signatures of industrialized and non-industrialized lifestyles. We identified 1,438 KEGG functions (21% of all KOs) whose marginal variation is more strongly associated with lifestyle than with diet or genetics, and that show consistent, significant associations with industrialization (PC1_Lifestyle) across countries (Fig. 3E). We then performed a KEGG pathway module enrichment analysis, identifying modules significantly enriched in either lifestyle category and composed of these cross-country, industrialization-associated KOs. We discovered that three different modules involved in cobalamin (vitamin B12) biosynthesis are enriched in the microbiome of industrialized societies (Fig. 3F). Interestingly, we recently found that the cob operon, which encodes enzymes for cobalamin biosynthesis, contains single-nucleotide variants convergently associated with industrialization across multiple gut bacterial species ^36^, suggesting that natural selection and adaptive processes are acting on this function in the gut microbiome. We also found that the propanoyl-CoA metabolism pathway (module M00741), which is involved in the metabolism of the short-chain fatty acid propionate ^37,38^, is enriched in industrialized societies (Fig. 3F). Intriguingly, the propanoyl-CoA pathway contains the methylmalonyl-CoA mutase (MUT) enzyme, which uses cobalamin as a cofactor (Fig. 3F). Looking more broadly at all known enzymes of gut bacteria that use cobalamin as cofactor (see Methods), we found a significant enrichment of multiple cobalamin-dependent enzymes in the microbiome of industrialized societies (Supp. Fig. 3D). This includes the lysine 5,6-aminomutase, which is used by opportunistic pathogens, such as *Fusobacterium nucleatum* – involved in colorectal cancer – to ferment lysine to produce butyrate, acetate, and ammonia ^39^. Further experiments will be needed to validate the increased cobalamin production in the microbiome of industrialized societies, and to understand the consequences of this increased cobalamin availability on microbiome functions and dysbiosis.

### Loss of gut microbiome stability and emergence of co-abundant taxa sets among populations living in industrialized societies

We next sought to determine whether lifestyle-associated shifts in the microbiome impact the structure of co-abundance networks and the stability of microbial communities. To explore this, we reconstructed species-level co-abundance networks based on microbial species profiles of industrialized and non-industrialized cohorts ^40^ (SparCC correlations, retaining correlations with |cor| > 0.3). We compared network characteristics while controlling for differences in total number of taxa, number of hosts, alpha diversity, age, sex, BMI and host geographic distribution between the two groups (see Methods, Supp. Fig. 4A-E and Supp. Table 4) (Fig. 4A). Our analyses revealed that the microbial network of industrialized individuals exhibit significantly higher node degree (Wilcoxon test; p-val = 8.2e-09; controlling for alpha diversity: p-val = 3.9e-05) (Fig. 4B & Supp. Fig. 4F-H). To exclude the possibility that this pattern reflected greater ecological heterogeneity in non-industrialized populations compared to the more homogeneous environments of industrialized groups, we reconstructed co-abundance networks within each locality (n >= 30 individuals) and ranked them along PC1 Lifestyle. We found a positive correlation between node degree and PC1 Lifestyle (R^2^ = 0.3, p-val = 8.7e-03; Fig. 4B), confirming that more urbanized and industrialized populations harbor microbial networks with higher connectivity. We further found that the network of non-industrialized individuals is enriched in negative correlations compared to the industrialized network (22% vs. 5% of all significant correlations with |rho| > 0.3 are negatives; Fisher Test, Odds ratio = 5.7, p-val = 7.4e-207; Fig. 4A). Overall, industrialized microbiomes are characterized by low alpha diversity (Fig. 2), high node degree, and an enrichment of positive interactions – features previously associated with reduced microbiome stability across environments ^41–44^. Consistent with this, we confirmed through direct stability measurements that industrialized microbiomes exhibit lower network stability (Fig. 4C). While lower community stability is found in many chronic diseases and linked to lower resistance to pathogen invasion ^45^, whether reduced microbial community stability of healthy industrialized populations increases their susceptibility to disease remains to be experimentally validated.

**Figure 4.**
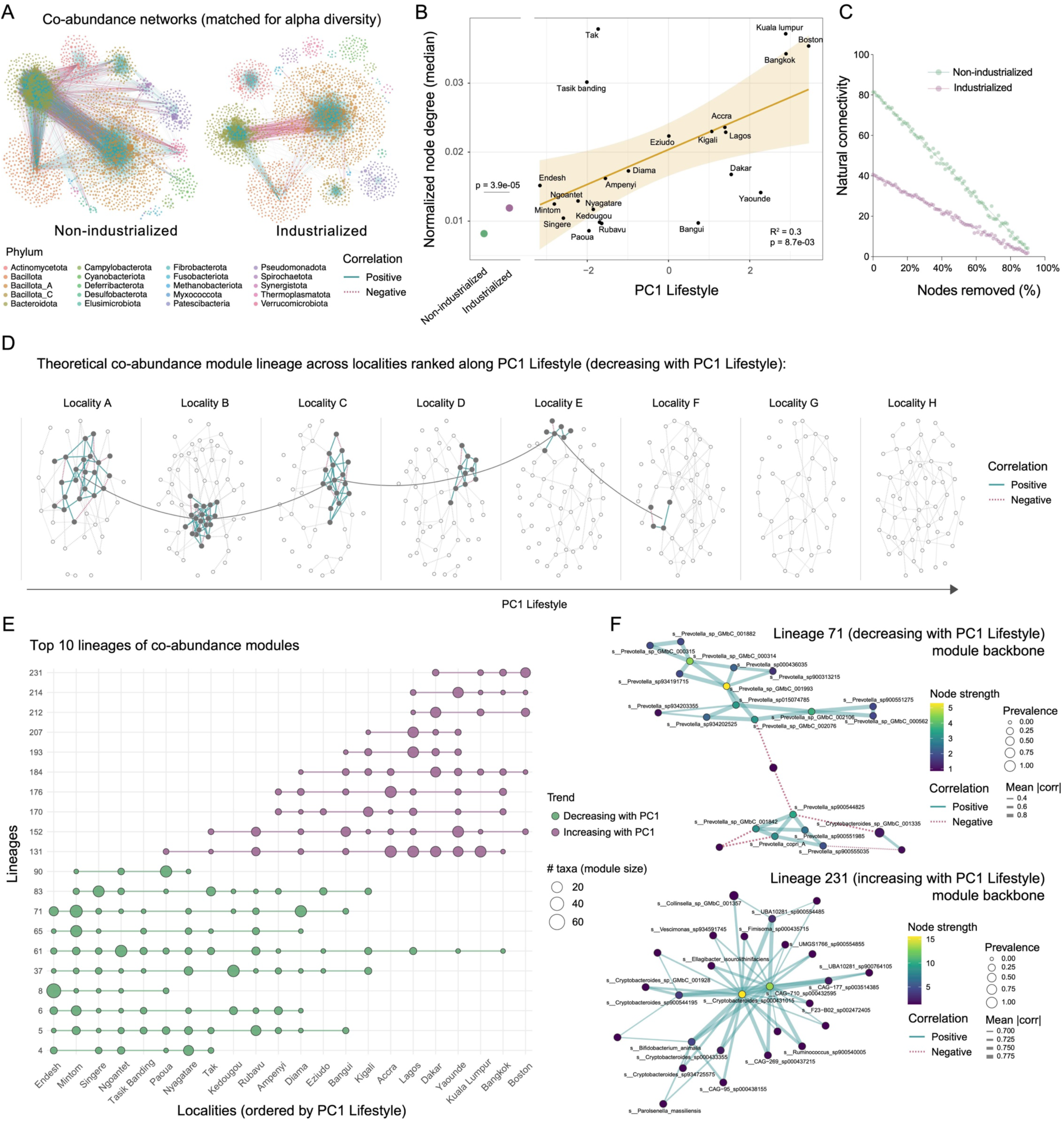
Decreased gut microbiome stability among populations living in industrialized societies. A. Co-abundance networks reconstructed from metagenomic profiles of non-industrialized (left) and industrialized (right) samples. Only significant compositionally-corrected abundance correlations with |coefficient| >= 0.3 are retained (edges). Each node represents a bacterial species. Co-abundance networks were reconstructed by matching the two groups for alpha diversity (n=460 individuals in each group, see Methods and Supp. Fig. 4). Networks controlling for age, sex, BMI and geographic distribution are shown in Supp. Fig. 4. B. Increased node degree among co-abundance networks of more industrialized populations. Normalized node degree was calculated at the aggregate level for both non-industrialized and industrialized groups (left panel, Wilcoxon test), as well as for each individual locality (n participants >= 30) (right panel, linear regression with PC1 Lifestyle, coefficients shown at the bottom right). Average position along PC1 Lifestyle was calculated for each locality. C. Natural connectivity measurement of co-abundance networks based on industrialization status. Nodes were progressively and randomly removed (x axis) to simulate perturbations and measure the stability of the networks. D. Toy schematic illustrating the concept of a lineage as a series of matched co-abundance modules tracked across ordered localities. Eight synthetic locality-specific co-abundance networks are shown for eight localities ordered by PC1 Lifestyle. The highlighted module (colored edges and filled nodes) depicts a lineage negatively associated with PC1 Lifestyle, with module size diminishing along the PC. Curved grey connectors link the centroids of the highlighted module. Edge color indicates correlation sign. E. Top-10 module lineages with positive/negative trends along PC1 Lifestyle. Localities (x-axis) are ordered by mean PC1 Lifestyle; lineages (y-axis) are labeled (see Supp. Table 4). Points show the presence and size of the module in a given locality. Modules were matched across adjacent localities using taxonomic compositional overlap (Szymkiewicz– Simpson), with one-to-one assignment (Hungarian) and a single-gap lookahead to accommodate for module absence. Colors indicate trend category: green = negatively associated with PC1 Lifestyle (markers of less industrialized contexts), purple = positively associated (markers of more industrialized contexts). Top 10 lineages per trend were selected by module size and persistence over localities. F. Backbone co-abundance networks for two representative lineages. For each lineage, we identify the largest module occurrence (peak size) and build a concise correlation backbone over this set of taxa. Node size reflects prevalence of the taxon across localities where the lineage is present; edges represent the mean pairwise correlation across those localities. To improve readability, loosely connected nodes were removed from the co-abundance backbone. Top: lineage negatively correlated with PC1 Lifestyle; bottom: lineage positively correlated with PC1 Lifestyle.

We then searched for modules of co-abundant taxa that are markers of host lifestyle and industrialization, using a multi-layer network approach (see Methods). We detected modules in all locality-specific networks, and matched them across localities ordered along PC1 Lifestyle to define module lineages (i.e. recurring co-abundant taxon sets tracked along variations in host lifestyle) (Fig 4D). For each lineage, we correlated module size per locality with PC1 Lifestyle and identified multiple lineages that are either positively (n = 38) or negatively (n = 72) associated with PC1 Lifestyle (Fig. 4E&F, Supp. Fig. 4I, Supp. Table 4). To assess whether these counts could arise by chance, we performed a label-swap permutation that randomly reassigns locality PC1 values. We found that observed counts are higher than null expectations (p = 0.002 and p < 0.001 for positively and negatively-associated lineages, respectively; Supp. Fig. 4J), indicating directional reorganization of co-abundance structure along host industrialization levels, with specific module lineages building up or fading as industrialization increases.

### Fecal biomarkers link immunity and inflammation with host lifestyle and microbiome composition

Next, we tested whether our observed shifts in microbiome composition and stability linked to host lifestyle (Fig. 2–4) are associated with host physiology and immunity. For this, we investigated fecal protein biomarkers across GMbC populations, and explored their association with the gut microbiome (Fig. 5A-E, Supp. Table 1). We measured fecal levels of secretory immunoglobulins A (sIgA) and M (sIgM) (Fig. 5B & E) as markers of mucosal immune responses against microbial antigens ^46^, chromogranin A (CgA) (Fig 5C) as a marker of gut stress and neuroendocrine system activation ^47^, and calprotectin (Fig 5D) as a marker of intestinal inflammation ^48^. We found that fecal IgA is positively correlated with both CgA and calprotectin, correcting for Age, Sex, BMI, Geography and Genetics (LM models, q-val = 1.9e-13 and q-val = 3.5e-04, respectively) (Supp. Fig. 5A). This confirms that elevated IgA secretion frequently co-occurs with gut inflammation and homeostatic stress ^46^. Yet, the substantial unexplained variance across biomarkers confirms that they reflect distinct aspects of gut physiology and immune activity.

**Figure 5.**
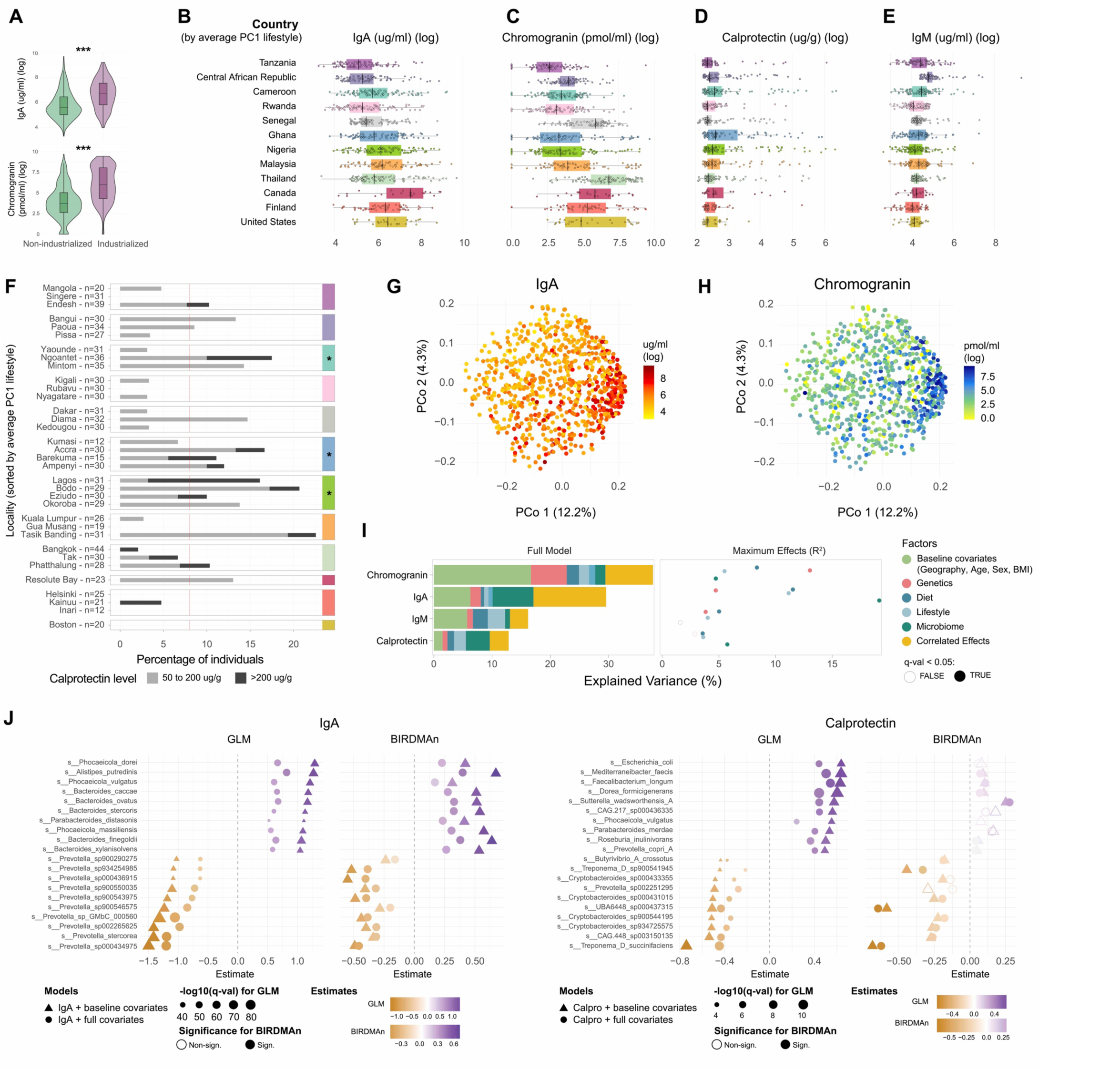
Industrialized and urban lifestyles are associated with elevated levels of fecal markers of inflammation and immune response. A. Fecal IgA and chromogranin levels based on industrialization status (Wilcoxon tests, p-val = 3.6e-18 and p-val = 4.2e-30, respectively). B-E. Distribution of fecal IgA, chromogranin, calprotectin and IgM levels across countries, sorted by average position along PC1 Lifestyle, in ascending order. F. Percentage of participants with intermediary (50 to 200 ug/g) or high (>200 ug/g) levels of fecal calprotectin, across localities. The red dash line indicates the mean percentage of participants with calprotectin > 50 ug/g across all localities. Countries with a significant enrichment of participants with moderate to high levels of calprotectin (> 50 ug/g) are shown with asterisks (logistic regression with a GLM and a binomial distribution; Cameroon: p-val = 0.04; Ghana: p-val = 0.02; Nigeria: p-val = 0.03). Countries are colored as in panels B-E. G-H. PCoA of species-level microbiome compositions (as in Fig. 2B), with participants colored by level of IgA and chromogranin. I. Variance partitioning of IgA, Chromogranin, Calprotectin and IgM levels. The same statistical framework as in Fig. 2 and 3 was used, adding PCs of microbiome compositions to the models (dark green) to account for and measure the contribution of the microbiome to fecal biomarkers. J. Identification of bacterial taxa associated with fecal IgA levels (left panel) and fecal calprotectin levels (right panel). Both GLM and Bayesian-based (BIRDMAn) approaches were used to detect taxa being consistently differentially abundant based on IgA or calprotectin. Two regression models were tested with each approach. In the first one, the model controls for baseline host covariates (age, sex, BMI and geography) (triangle symbols). In the second model, host Lifestyle, Diet and Genetic PCs are added to further control for the effect of host covariates (circle symbols). Top 10 bacteria with the strongest positive and negative effect sizes based on GLM runs are represented (See Supp. Table 5 for the full data).

Our analysis shows elevated sIgA levels in industrialized populations (e.g., US, Canada, Finland) compared to non-industrialized ones (mean sIgA = 772 vs. 327 µg/mL; Wilcoxon p = 3.6e-18), and in urban vs. rural individuals (mean sIgA = 620 vs. 299 µg/mL; Wilcoxon p = 3.1e-21) (Fig. 5A&B). This trend is confirmed at higher resolution within most countries, with higher sIgA in urban compared to rural populations (Supp. Fig. 5B). Adjusting for demographic, geographic, and genetic factors confirmed these findings (GLM; industrialization p = 2.5e-05; urbanism p = 2.7e-12). This is further confirmed when looking at populations with the same self-declared ethnicity having different urbanism status, such as Beti individuals in Cameroon, who exhibit higher fecal IgA levels in Yaounde (urban) compared to Ngoantet (rural) (Wilcoxon test, p-val = 0.0003). Intestinal eukaryotic microbes may elicit specific sIgA responses and contribute to fecal IgA levels ^49^. Yet, we found that non-industrialized groups have higher prevalence of intestinal eukaryotic microbes (Supp. Fig. 5C), while exhibiting lower fecal IgA levels (Fig. 5A). We also did not find strong associations between sIgA and levels of Entamoeba or Blastocystis (Supp. Fig. 5D & Supp. Table 5). We next tested for a potential compensatory IgM secretion in non-industrialized regions, as observed in IgA-deficient patients ^50^. We found no signal for such compensation effect, as IgA and IgM are positively correlated both at the aggregate level (Spearman rho = 0.13, p = 3.4e-05) and among non-industrialized regions (rho = 0.21, p = 2.3e-09). Finally, we verified whether sIgA levels in non-industrialized groups could be explained by a higher abundance of IgA proteases in the metagenome ^51^. We observed the reverse trend, with lower levels of proteases in non-industrialized groups (Pearson correlation; beta = 0.74, p = 1.57e-07) (Supp. Fig. 5E).

Next, we hypothesized that differences in sIgA levels are related to differences in the microbiome. We found that sIgA levels are strongly explained by variations in microbiome composition at the aggregate level (PERMANOVA; IgA: R² = 0.06, p < 0.001; Fig. 5G), as well as within each country (Supp Fig. 5F). We partitioned sIgA variance using the same approach as above, with the addition of the microbiome as a predictor in the model (see Methods) (Fig. 5I & Supp. Table 5). We found that the microbiome has the highest maximum effect on sIgA (R2 = 0.17, p-val = 4.4e-35), and the highest marginal effect compared to Lifestyle, Genetics and Diet (R2 = 0.07, p-val = 3.6e-15) (Fig. 5I). Using a differential abundance analysis ^52^, we further identified individual bacterial taxa that are associated with sIgA levels (Fig. 5J & Supp. Table 5), with Phocaeicola/Bacteroides and Prevotella SGBs being positively and negatively associated with sIgA, respectively.

Variation in fecal CgA shows contributions of Lifestyle and Genetics, and, to a lesser degree, the microbiome. We observed that (i) CgA is elevated in individuals of industrialized and urban lifestyles compared to non-industrialized and rural lifestyles, respectively (mean CgA = 412 vs. 48 pmol/ml; Wilcoxon p = 4.2e-30; mean CgA = 153 vs. 48 pmol/ml; Wilcoxon p = 5.5e-14) (Fig. 5A&C & Supp. Fig. 5G); (ii) CgA is associated with community-level microbiome variation (PERMANOVA; CgA: R² = 0.05, p < 0.001; Fig. 5H); and (iii) variance partitioning reveals that Genetics is a strong driver of CgA (maximum R2 = 0.13, p = 1.3e-25; marginal R2 = 0.06, p = 2.5e-12), while the microbiome has a lower contribution (maximum R2 = 0.05, p = 2.1e-09; marginal R2 = 0.02; p = 4.5e-04) (Fig. 5I). As for sIgA, we did not find strong associations between CgA and eukaryotic microbes, to the exception of a weak negative association with Blastocystis (LM model, beta = -0.3, p = 0.04, Supp. Fig. 5D, Supp. Table 5).

Looking at calprotectin, we found that 8.9% (n=85) of GMbC participants had mild or active intestinal inflammation (> 50ug/g stool). Among those, 21% individuals (n=18) had active inflammation (> 200ug/g stool). We did not find a significant correlation between calprotectin and industrialization or urbanisation status at the cohort level. However, we observed an increased frequency of moderate-to-high calprotectin in sub-Saharan Africa, irrespective of urbanization status, including Nigeria (p = 0.03), Cameroon (p = 0.04) and Ghana (p = 0.02) (Fig. 5F). We also observed a higher frequency of moderate inflammation in rural, low-density populations that were recruited in Malaysia (Tasik Bandig) and Thailand (Phatthalung) (Fig. 5F). These groups are currently facing rapid lifestyle, dietary and environmental perturbations, which we witnessed during our visits, such as resettlements outside their original land ^53,54^. We found a significant association between calprotectin levels and microbiome compositions (PERMANOVA, R² = 0.01, p < 0.001). Individuals with elevated calprotectin tended to have microbiome profiles intermediate between non-industrialized rural and urban industrialized populations, consistent with the intermediary compositions observed in sub-Saharan countries (Fig. 2B & Supp. Fig. 5H).

At the taxon level, effect sizes were generally smaller than for sIgA (Fig. 5J & Supp. Table 5), but we observed a consistent negative association with the short-chain-fatty-acid producer *Treponema succinifaciens* ^55^, which tends to be depleted in industrialized and urban populations (Fig. 3B). Interestingly, using microbiome-colonocyte co-culture assays, we recently found that Treponema is associated with the expression of hypoxia and TNF signaling genes, which are involved in cellular immune modulation by bacteria ^36^. This suggests that elevated inflammation may contribute to the depletion of *Treponema succinifaciens*, positioning it as a potential marker of healthy microbiomes. We also tested for associations between calprotectin and known pathobiont species (n = 100 species, see Supp. Table 5). Among those, calprotectin was positively associated with three enteric pathobionts (*Escherichia coli*, *Enterococcus faecalis* and *Klebsiella pneumoniae*, Supp. Fig. 5I). Localities with excess of high-calprotectin individuals (Fig. 5F) are partially characterized by the presence of these pathobionts in the metagenomes, with many cases of co-occurrence (presence of ≥2 pathobionts) (Supp. Fig. 5I). Because these three taxa are also co-enriched in inflammation, such as inflammatory bowel disease (IBD) ^56^, the elevated calprotectin and microbial dysbiosis may be explained by either ongoing pathogenic infection or intestinal inflammation.

Overall, these findings demonstrate that the gut microbiome of industrialized and recently urbanized or settled populations is associated with increased IgA secretion, homeostatic stress, and gut inflammation. These results underscore the need for future epidemiological studies to evaluate the prevalence of intestinal inflammation and dysbiosis in these populations (*e.g.* across sub-Saharan African metropolitan areas), in order to guide potential intervention initiatives.

### Industrialization is linked to shifts in IgA-gut bacteria interactions

Having established that the level of fecal IgA is a marker of host lifestyle and is strongly associated with the microbiome (Fig. 5), we next sought to investigate whether IgA-gut bacteria interactions change across human populations and reflect host lifestyle. For this, we sorted, identified and quantified gut bacteria bound by IgA, using IgA-Sequencing (IgA-Seq) ^46,50^ (see Methods). We calculated an IgA coating index (ICI) to quantify how strongly individual bacteria are coated by IgA (see Methods). We applied IgA-Seq on a subset of 300 individuals of the GMbC cohort covering broad geographies and lifestyles, retaining 283 samples for downstream analysis after post-sequencing rarefaction and filtering (see Methods).

We first tested whether IgA-bacteria interactions are evolutionarily conserved in humans. We found a strong correlation between the average ICI of taxa occurring in both industrialized and non-industrialized groups (Pearson r = 0.73, p = 2.2e-29; Fig. 6A, adjusted for sample size differences; see Methods). We observed the same trend when considering prevalence of coated bacteria (i.e., frequency of individuals with ICI ≥10 ^46^, r = 0.67, p-val = 0.003, Supp. Fig. 6A). We further examined key bacterial taxa previously shown to be highly coated by IgA in the gut: Akkermansia ^50,57^, Enterobacteriaceae ^50,58–60^, and *Bacteroides fragilis* ^61^. For these lineages, we found consistent patterns between the groups in average ICI (Fig. 6B) and IgA-coating prevalence (Supp. Fig. 6A). Overall, these results suggest that IgA-bacteria interactions are broadly conserved across human populations, despite major inter-population variability in the microbiome driven by shifts in lifestyle (Fig. 2 & 3). It also points to the presence of similar antigen profiles across gut bacterial strains, and common IgA induction mechanisms across human populations.

**Figure 6.**
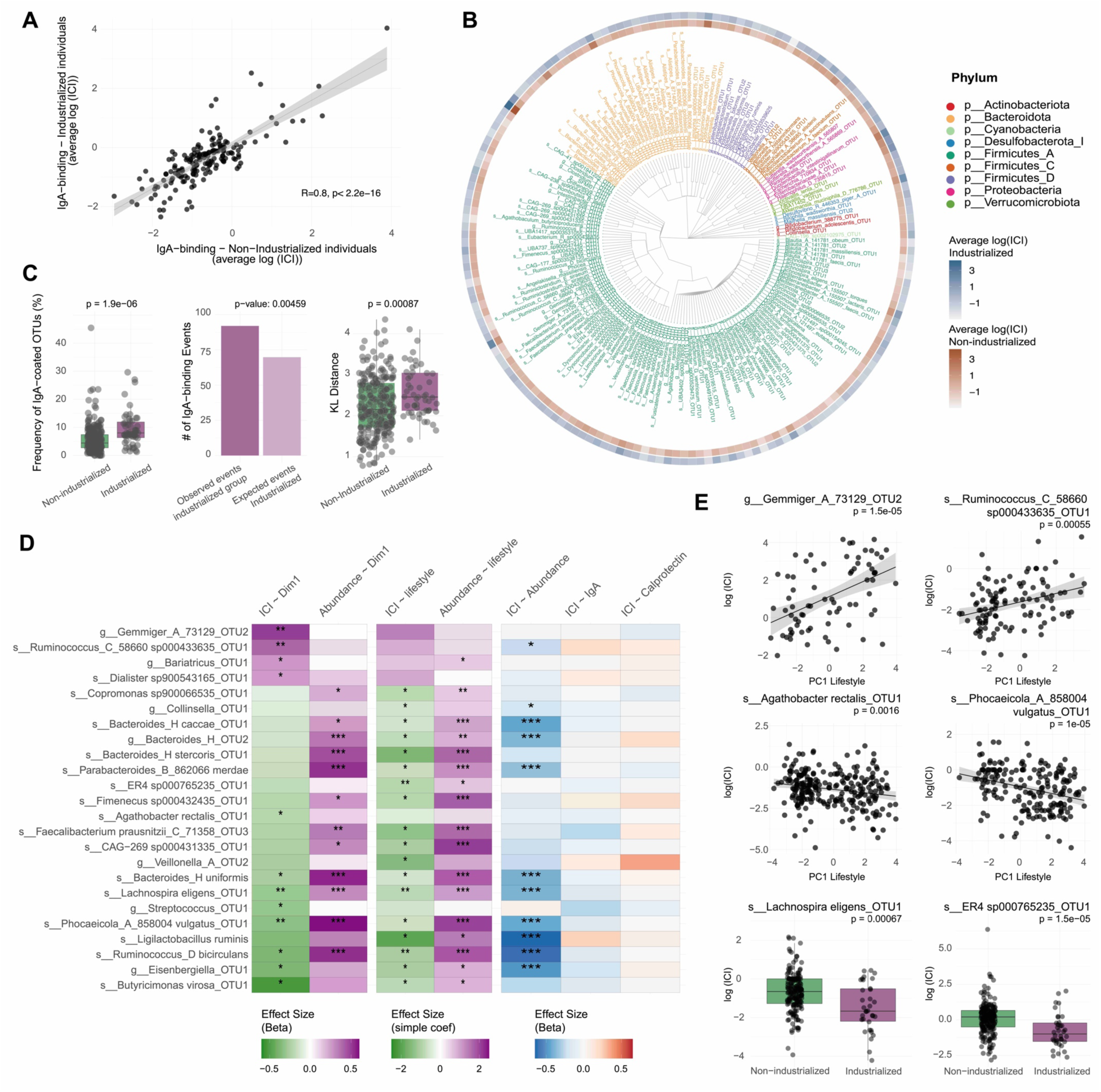
Shifts in IgA-gut bacteria interactions in the gut microbiome of industrialized populations. A. Correlation of OTU-specific average IgA Coating Index (ICI) between individuals from industrialized and non-industrialized groups (Pearson correlation of log ICI values). Comparisons were made on OTUs with sufficient representation and balanced sample size between lifestyle categories (n = 166, see Methods). B. Phylogenetic distribution of average ICI values in industrialized and non-industrialized groups (blue and red color gradients, respectively). OTUs are colored by phylum. C. Comparison of IgA-coating between industrialized (n = 51 samples) and non-industrialized (n = 232 samples) groups, based on: the frequency of IgA-coated OTUs (ICI > 10) per individuals (left); the comparison of observed vs. expected IgA-coating events in the industrialized group, based on the frequency of IgA-coating events in the non-industrialized group (p-value calculated with the Poisson distribution) (middle); and the Kullback–Leibler (KL) divergence between the IgA+ and the total 16S relative abundance profiles (right). D. Correlation between IgA coating index or relative abundance and industrialization status of the host. Compositionally-corrected (CLR-transformed) abundance values are used. OTUs with sufficient representation across lifestyles were included in the analysis (n = 197). Correlations between ICI and industrialization were calculated with both PC1_Lifestyle (ICI ∼ PC1_Lifestyle) and industrialization as a binary variable (ICI ∼ industrialized/non-industrialized). The correlation between the CLR-corrected abundance of OTUs and industrialization were also measured (Abundance ∼ PC1_Lifestyle & Abundance ∼ industrialized /non-industrialized). The correlation between ICI and relative abundance of the OTUs is also shown (ICI ∼ Abundance). *: p-val < 0.05; **: p-val < 0.01; ***: p-val < 0.001. E. Significant associations between ICI and industrialization shown for six different OTUs.

Despite broad conservation, we found multiple associations between IgA-coating and host lifestyle. First, the total frequency of highly coated bacteria (ICI ≥10) is higher in industrialized individuals than in non-industrialized individuals (Fig. 6C, Wilcoxon test, p = 1.9e-06; Poisson test, p = 0.004 - See Methods). This trend is also confirmed when comparing urban vs. rural individuals (Supp. Fig. 6B, genus-level data, Wilcoxon, p = 0.0043) and when correlating ICI with PC1_lifestyle (Spearman rho = 0.14, p = 0.025). Comparing the IgA+ fraction abundance profiles to the pre-sorted fraction using Kullback–Leibler (KL) divergence, we found higher KL divergence in industrialized and urban individuals (Fig. 6C, Wilcoxon tests, p = 8.7e-04 and p = 7.4e-05, respectively), indicating stronger IgA coating intensity in industrialized and urban individuals than expected based on background microbiome profiles, compared to non-industrialized and rural individuals.

Second, we identified various bacterial lineages with IgA coating patterns that are linked to host lifestyle. For example, *Phocaeicola vulgatus*, *Lachnospira eligens*, *Ruminococcus_D bicirculans*, and *Butyricimonas virosa* OTUs show an elevated IgA coating index in the non-industrialized category (Fig. 6D & E), while OTUs of Ruminococcus_C and Gemmiger have increased IgA coating in the industrialized category (Fig. 6D & E). The same Gemmiger OTU, along with *Akkermansia muciniphila*, have an elevated frequency of highly coated bacteria (ICI ≥ 10) in industrialized individuals (Supp. Fig 6C). In many cases, IgA coating and lifestyle are independently associated with bacterial abundance (Fig. 6D); for instance, *P. vulgatus* has both higher IgA coating and a lower abundance in the non-industrialized microbiomes. However, other taxa whose IgA coating profiles are associated with industrialization, such as Ruminococcus_C_sp000433635 and ER4_sp000765235, do not show a significant correlation between industrialization and their abundance in the microbiome. Overall, this suggests that a complex interplay between microbial community structure, host lifestyle, and immune responses contributes to shaping IgA coating, and that some bacterial strains may elicit host lifestyle-specific IgA responses independently of their abundance in the gut.

### Low portability of microbiome-based predictors of health and disease

Finally, we hypothesized that previously developed microbiome-based predictors of disease may have limited applicability to non-industrialized populations. This would mirror a well-recognized issue with polygenic risk scores (PRSs), which predict disease risk from human genetic variant frequencies. PRSs often fail to transfer across populations because they are typically trained on European ancestry cohorts and capture population-specific allele frequencies ^62^. Consistent with this limitation, we confirmed that an IBD PRS derived from European cohorts ^63^ showed low predictive accuracy when applied to non-European GMbC participants (Methods, Supp. Fig. 7A-B). We then tested our hypothesis with a microbiome-based predictor of health on the GMbC metagenomic data. For this, we used the gut microbiota health index (GMHI), which accounts for the abundance of “health” and “disease” marker taxa ^64^. Developed from microbiome data of industrialized populations, the GMHI effectively classified most industrialized GMbC individuals as healthy (Fig. 7A). In contrast, it performed poorly on metagenomes of non-industrialized societies, owing to the lower prevalence of GMHI marker taxa in the microbiome of these populations (Fig. 7A). This pattern held across both country and admixture groupings (Fig. 7B&C).

**Figure 7.**
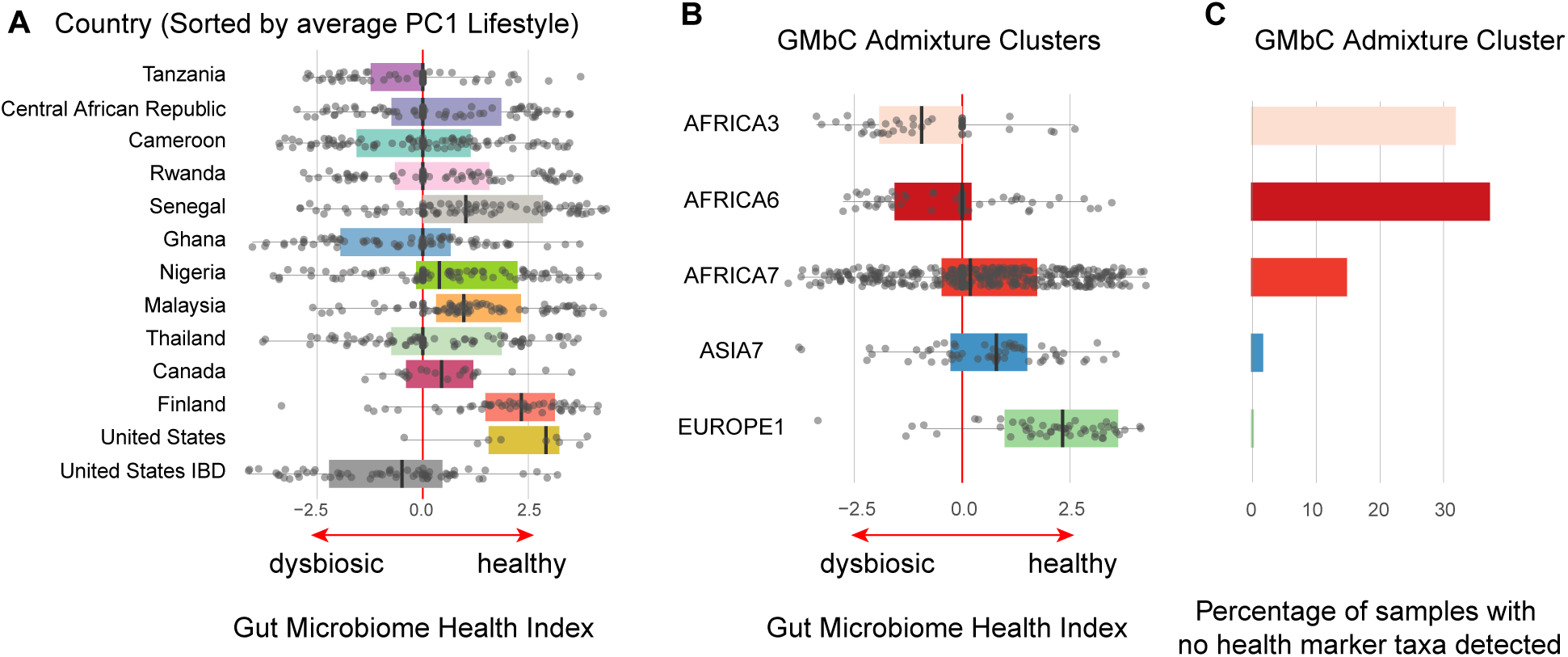
Low portability of microbiome-based biomarkers of health and disease. A. Distribution of Gut Microbiome Health Index (GMHI) across GMbC countries. GMbC participants do not have known chronic diseases. The GMHI was also calculated on an external validation cohort of inflammatory bowel disease (IBD) patients from the USA (in grey; see Methods). Median GMHI values are shown as black lines. Positive values indicate healthy microbiomes, while negative values indicate dysbiotic microbiomes. Median values for Tanzania, CAR, Cameroon, Rwanda, Ghana and Thailand is 0, suggesting an absence of information to call for healthy or dysbiotic microbiomes. B. GMHI values across admixture clusters. Admixture clusters with a minimum of 50 individuals were included in the analysis. C. Percentage of samples that do not contain any of the health marker taxa that are informative for the GMHI, across admixture clusters. The lack of GMHI accuracy observed in panels A and B is due to the absence of key maker taxa of health defined from cohorts of majority populations.

## Discussion and limitations

Our study establishes a robust framework for dissecting the specific contributions of individual factors to gut microbiome variability across diverse global populations. Yet, we acknowledge that a significant portion of variation is still attributed to the collinearity between factors (“non-additive effects”, Fig. 2 & 3). Some effects may never be fully dissected, owing to the strong association between, e.g. lifestyle and diet. In addition, the underperformance of microbiome-based disease predictors in populations of non-European descent reveals the need to incorporate clinical cohorts from low- and middle-income countries of microbiome-associated chronic diseases that are commonly studied in majority countries, such as IBD or Type 2 diabetes. Understanding the diversity of microbiome shifts that occur in these diseases across worldwide regions will help building markers that are tailored to specific groups, allowing them to benefit from improved early risk prediction, prevention, and personalized medicine. The future inclusion of additional worldwide populations using the GMbC framework, including such clinical cohorts, will increase our capacity to tease apart and leverage the effects that various host factors have on the gut microbiome.

Our findings revealed increased levels of gut stress markers among healthy populations of industrialized and urban cohorts that are associated with the gut microbiome. Whether the microbiome occurring with industrialization and urbanization is more prone to dysbiosis, contributes to perturbations of host physiology, or increases vulnerability to infections ^15,65^ remains to be experimentally tested. In addition, understanding which specific attributes of the industrialized microbiome (e.g. lower diversity, decreased stability, or specific microbes) drive non-healthy host phenotypes will require further investigation. Interestingly, we recently used co-culture experiments with GMbC fecal samples from various geographies and industrialization status and colonic epithelial cells to examine host transcriptional responses to microbiome variation ^36^. We found that microbiomes with lower diversity tend to elicit expression of genes involved in epithelial restructuring, while microbiomes of higher diversity triggered stronger host transcriptional responses overall. We further found that microbiomes of urban hosts promoted stronger transcriptional responses of genes involved in innate immune pathways, including TNF signaling and bacterial antigen recognition ^36^. Together with our profiling of inflammatory markers and IgA induction/recognition patterns, these results provide causal evidence that diversity and compositional shifts in the microbiome occurring with host industrialization and urbanization exert strong impact on host physiology. The GMbC biobank of preserved stool samples and microbiomes will serve as a critical resource for extending such mechanistic investigations in the future.

Moving forward, the GMbC aims to serve as a global observatory for continuous monitoring and understanding of changes in the human metaorganism that occur along the spectrum of lifestyle transitions. Through longitudinal and multimodal data collection across a range of populations and disease contexts, these efforts will support the development of *in situ* strategies for microbiome conservation and context-specific interventions. In doing so, the GMbC seeks to ensure that microbiome science is not only inclusive and globally representative, but also directly relevant to local populations by enabling local science ^66,67^, efficient responses to emerging health challenges ^68,69^, and informed decisions on trajectories of lifestyle transitions ^26^.

## Methods

### Participant recruitment and collection of biospecimens

We recruited 1,015 healthy adult participants across 12 countries. Individuals reported no symptoms of infectious or chronic disease at the time of enrollment. The selection of sampling locations and human populations within each country was done collaboratively by consortium scientists, with the goal of capturing environmental, lifestyle, and cultural diversity within each national context. Written informed consent was obtained from all participants, using translations in local language when appropriate. Research & ethics approvals were obtained from the MIT IRB (protocol #1612797956) and from the Ethics commission of the Medical Faculty of Kiel University (Studie D 511/24). Permits were also obtained in each sampled country prior to the start of sample collection, from the following ethics committees:

- Cameroon: Comité National d’Ethique de la Recherche pour la Santé Humaine, protocol #2017/05/901/CE/CNERSH/SP;
- Canada: Nunavut Research Institute, protocol #0205217N-M;
- Central African Republic: Comité d’Éthique et Scientifique de la Faculté des Sciences de la Santé de l’Université de Bangui, No. 17/UB/FACSS/CES20;
- Finland: Coordinating Ethics Committee of Helsinki and Uusimaa Hospital District, protocol #1527/2017;
- Ghana: Cape Coast Teaching Hospital Ethical Review Committee, protocol #CCTHERC/RS/EC/2016/3 and Committee on Human Research, Publication and Ethics of the Komfo Anokye Teaching Hospital, protocol #CHRPE/AP/398/18;
- Malaysia: Universiti Malaya Medical Research Ethics Committee, MREC ID No.: 2018219-6033;
- Nigeria: National Health Research Ethics Committee of Nigeria, protocol #NHREC/01/01/2007-29/04/2018.
- Rwanda: National Ethics Committee, protocol IRB 00001497 of IORG0001100
- Senegal: Comité National d’Ethique pour la recherche en Santé, No.: 0000073 MSAS/DPRS/CNERS
- Tanzania: National Institute for Medical Research, protocol #NIMR/HQ/R.8a/Vol. IX/2657;
- Thailand: The Human Research Ethics Committee of the Faculty of Medicine of Thammasat University No.1, Certificate of Approval 143/2019
- United States of America: Chief Dull Knife College, protocol #FWA00020985;

Participants self-collected a fresh fecal sample using sterile containers, which were returned to GMbC scientists for on-site processing within 0 to 3 hours following defecation. Raw stool was diluted 1:5 in a 25% pre-reduced anaerobic glycerol solution containing acid-washed glass beads for homogenization, then aliquoted into 2 mL cryogenic tubes. Aliquots in cryoprotectant were flash-frozen either in liquid nitrogen (−196 °C) using a cryoshipper tank or stored at −80 °C. An additional 1–2 g of stool was preserved in RNAlater for DNA stabilization. Saliva samples (2mL) were collected using the Oragene OGR-500 kit (DNA Genotek), and preserved at room temperature. All fecal and saliva samples were subsequently shipped to MIT for further processing and storage.

### Metadata collection

We collected metadata on lifestyle, environmental exposure, and medication use from all participants. A full list of metadata variables is provided in Supp. Table 1.

To determine the industrialization status of sampled populations, we used the Human Development Index (HDI), as used previously ^25^. HDI values at national and sub-national levels were obtained from the Global Data Lab database (https://globaldatalab.org/). When sub-national data were unavailable for a given locality, the national-level HDI was used. Populations from localities with HDI values below the national median (0.739 in 2022) were classified as non-industrialized, while those with values above this threshold were classified as industrialized.

To assess the urbanization status of each community, we estimated population density using the SEDAC Population Estimation Service, based on GPS coordinates (https://sedac.ciesin.columbia.edu/mapping/popest/pes-v3/). We estimated the population size within a 5 km radius of each locality and calculated population density as the number of inhabitants per square kilometer of land area. Populations with densities above 1,000 inhabitants/km² were classified as urban, and those below as rural.

We also report the subsistence strategy of each population, defined as the primary means by which inhabitants obtain essential nutrition for survival. Populations were classified into eight broad subsistence categories: Foraging, Foraging & Farming, Farming, Pastoralism, Fishing, Foraging & Industrialism, Farming & Industrialism, and Industrialism. Detailed definitions of these categories are provided in Supplement Information, Table 1. Classifications were based on direct field observations and interviews with local stakeholders. Subsistence strategies are inherently dynamic, often mixed, and shaped by seasonal, ecological, and economic factors. We acknowledge the limitations of assigning populations to discrete categories and therefore treat these classifications as part of a broader framework. To better capture the complexity of human lifestyles, we integrated subsistence strategy data with other societal and environmental variables in dimensionality reduction analyses (see below), allowing individuals and populations to be represented along continuous lifestyle variables.

We collected data on self-declared ethnicity from each participant (Supp. Table 1). Many ethnic labels worldwide carry a complex history, sometimes originating from colonial-era classifications. Our interactions with every community made us also understand that they may choose themselves to reclaim these names as a source of identity, representation, and collective advocacy. For this reason, all our ethnicity data are explicitly recorded as “self-declared”, to represent the own identification of each participant. For a few participants, multiple self-declared ethnicities were recorded. In addition, we chose to perform our analyses of microbiome variations on genotyping data alone rather than on ethnic group labels, ensuring that observed microbiome patterns are driven by underlying genetic diversity rather than by potentially arbitrary or anachronistic ethnic groupings.

We collected dietary data using food frequency questionnaires (FFQs) tailored to each population. In every country, we adapted a standardized core questionnaire by incorporating additional food items that were relevant to the local diet. Questionnaires were then retrospectively updated to ensure consistency and comparability across populations while capturing dietary specificity.

We collected data on recent medication consumption, such as antibiotics and antiparasites (Supp. Table 1). Recent antibiotic/antiparasite consumption is rare in our cohort, limiting statistical power to detect a significant contribution of these exposures to microbiome variations (see Supp. Information).

We report climate data (annual mean temperature, temperature seasonality, annual precipitation and precipitation seasonality) of the year of sampling for each community in Supp. Table 1. Briefly, we downloaded historical WorldClim ^70^ climate data at 10mins resolution from https://gitlab.com/tpoisot/BioClim/tree/master/assets (wc2.0_bio_10m_precipitationSeason.tif, wc2.0_bio_10m_annualprecipitation.tif, wc2.0_bio_10m_tempSeasonality.tif and wc2.0_bio_10m_annualtemperature.tif), and estimated climate values at the year of sampling for each location using the QGIS tool.

### Dimensionality reduction of Lifestyle and Diet metadata

The following host metadata factors were categorized as “Lifestyle” variables: “locality_density” (# of inhabitants per km/sq), “industrialization_status” (industrialized: high HDI; non_industrialized: low HDI), “subsistence_strategy”, “water_source”, “c_section”, “breast_fed”, “household_size”, “animal_contact”, “tobacco”, “exercice_per_week”, “access_to_electricity”, “floor_type”, “travel_abroad_lastYear”, and “purge”. See Supp. Table 1 for a full description of these factors and their values. Missing data (representing only 3% of fields across all these factors) were imputed using the imputeFAMD function from the missMDA R package with 8 components. The multivariate analysis of lifestyle factors was then conducted with the PCAmix function from the PCAmixdata R package. PCAmix internally performs normalization of variables. We calculated the first 10 components for Lifestyle and Diet.

We used a previously developed tree-based method for analysis of food data on our FFQ profiles ^71^ (github.com/knights-lab/Food_Tree). Briefly, this method uses a hierarchical tree of food items to calculate UniFrac beta diversity dissimilarities of food profiles, and was shown to produce dietary distances that efficiently captures gut microbiome variations ^71^. Following the initial methodology based on the FNDDS Food Code and the ASA24 database, we manually added food items of our FFQ that were missing from the backbone topology of food items. We then used this food tree topology and the GUniFrac package to calculate unweighted and weighted UniFrac dietary distances between individuals (Supp. Table 1). Missing FFQ data were imputed with the imputePCA function of the missMDA package with 8 components. We then used the *cmdscale* function of the R stats package to calculate a PCoA, with Cailliez correction. The food tree used in this analysis is provided in Supp. Table 1.

To measure colinearity of host factors with PC1 Lifestyle, we measured the Variance Inflation Factor (VIF) index for each factor (vif function in the ‘car’ R package), which indicates how much the variance of a regression coefficient is inflated due to multicollinearity. A higher VIF indicates more collinearity. VIF values are reported in Supp. Table 1. Typically, a VIF > 5 suggests high collinearity.

### Shotgun metagenomic sequencing

DNA extraction and library preparation were performed following the same protocols as in our previous work ^25^. Briefly, stool samples stored in RNAlater were used for DNA extraction, using the MoBio Powersoil 96 kit (now Qiagen Cat No./ID: 12955-4). Genomic DNA libraries were constructed from 1.2ng of cleaned DNA using the Nextera XT DNA Library Preparation kit (Illumina) according to the manufacturer’s recommended protocol, with reaction volumes scaled accordingly. Before sequencing, libraries were pooled in equal quantities into multiple batches. Insert sizes and concentrations of each pooled library were determined using an Agilent Bioanalyzer DNA 1000 kit (Agilent Technologies). Paired-end sequencing (2×150-bp reads) was performed on a NovaSeq S4 platform (Illumina Inc) at the Broad Institute, yielding an average of 25.5 million reads and a median of 21.7 million reads per sample.

### Human gut bacterial isolate genomes from the GMbC and BIO-ML collections

Genomes of human gut bacterial cultured isolates were added to MAGs reconstructed from the GMbC metagenomes (see below) to build the reference SGBs used for taxonomic profiling. Isolate genomes that we generated in previous studies from both the BIO-ML ^72^ (n = 3,632) and GMbC ^25^ (n = 4,149) collections were considered. We also recently generated an additional set of 1,851 isolate genomes from GMbC participants, bringing the total number of isolate genomes used to reconstruct reference SGBs to 9,632 ^73^. All isolate genomes were cultured and sequenced using the same procedures and protocols ^25,72^.

### MAG assembly and SGB reconstruction from MAGs and isolate genomes

Short read data was assembled into contigs using MEGAHIT ^74^. We then employed four binning algorithms (MaxBin2 ^75^, MetaBAT 2 ^76^, CONCOCT ^77^ and VAMB ^78^) to reconstruct multiple sets of contig bins. Outputs were subsequently merged and refined using MAGScoT ^79^. For the initial binning, preprocessed sequencing reads of each sample were mapped against their contigs (minimum size >= 2kbp) using minimap2 (v2.24-r1122) and the short-read preset. The resulting SAM files were converted to sorted binary format (BAM) with samtools (v1.17). BAM files were used to estimate per-contig coverage using the jgi_summarize_bam_contig_depths script as input for binning in Metabat2 (v2:2.15) and MaxBin2 (v2.2.4). For CONCOCT (v1.1.0), as recommended by the developers, contigs were cut into chunks of 10kbp and used together with the BAM files for the binning procedure. While the previous binning tools were individually run on each sample, VAMB binning was performed on sets of multiple samples based on geographic sampling location. For this, length-filtered contigs were merged into a single fasta file which was used as mapping reference for minimap2. Per-contig coverages were estimated using the jgi_summarize_bam_contig_depths script, and then used in the main VAMB algorithm. The GTDB-defined set of 120 bacterial and 53 archaeal single-copy marker genes (r207) were used as reference for MAGScoT to find the highest scoring set of MAGs from the set of four binning tools per sample. MAGs with contamination levels higher than 10% and quality scores lower than 0.5 were filtered out, yielding a total of 24,163 MAGs (See Supp. Table 1 for MAG assembly statistics).

Next, we clustered MAGs and isolate genomes from the GMbC and BIO-ML isolate genome collections into species-level genome bins (SGBs). For this, we employed an iterative clustering workflow that maximizes quality of SGBs. We first performed an initial clustering of MAGs separately within each sampled geographic location at 97% similarity using dRep (v3.4.0) and average nucleotide identity (ANI). We selected a MAG representative within each cluster, selecting the MAG with the highest quality score. We classified MAG clusters into high quality (HQ; score > 0.7) and medium quality (MQ; 0.5 > score >= 0.7) clusters. As less complete genomes are more likely to form spurious genome clusters, only HQ cluster representatives from all geographic locations went into further clustering. HQ MAG representatives, along with GMbC and BIO-ML isolate genomes, were then clustered at 95% ANI SGBs, again selecting the highest scoring genome as HQ SGB representative. Previous MQ MAG representatives were then compared to the set of HQ SGBs using FastANI (v1.33) and assigned to an SGB cluster in case of >= 95% similarity. MQ clusters without a hit at >= 95% similarity with SGB representatives were clustered together at 95% similarity using dRep to form MQ SGBs. These SGB clusters were only retained if they consisted of more than a single genome at either the location-based pre-clustering step, or at the final MQ clustering step. HQ and MQ SGB clusters were then merged to form the final set of non-redundant SGBs (n = 434).

### Taxonomic abundance profiles and diversity measurements

The highest-scoring SGB representative genomes were used to assign taxonomic annotations to the SGB clusters using the GTDB-Tk (v2.3) and GTDB reference database (rel214) ^80^. In addition, the SGB representatives were used as reference genomes for taxonomic abundance estimation using salmon (v0.8.1) in metagenome mode (--meta) ^81,82^. The resulting per-contig abundances were further processed to obtain per-SGB relative abundances in analogy to TPMs (transcripts per million mapped reads). To minimize spurious detections, SGBs with fewer than 1,000 total reads and less than 250 TPMs (<0.025% relative abundance) were discarded. SGB-level abundances and taxonomic labels were then used to calculate taxon abundances from species to domain level by summing TPM values. To control for compositionality, TPM data were CLR-transformed using the clr function from the “compositions” R package, adding a pseudo count of 1 to the TPM abundances, and setting all post-transformation negative values to 0. The following functions and R packages were used to calculate alpha and beta diversity metrics – Faith’s PD: *pd* from the picante package; Shannon index: *diversity* from the vegan package; UniFrac distances: *GUniFrac* from the GUniFrac package; Aitchison: *dist* from the stats package. PERMANOVA tests were calculated with the adonis2 function from the vegan R package.

### Metagenome-derived gene cluster reconstruction, abundance profiling and annotation

We used Prodigal in metagenomics mode (-meta) to call for open reading frames on assembled contigs. We retained only full-length genes using the -c option. We then built sets of non-redundant 50% similarity gene clusters with MMseqs2 (version 10-6d92c) ^83^, using the easy-linclust workflow and the following options: --cov-mode 1 -c 0.8 --kmer-per-seq 80 --min-seq-id 0.50. Short read metagenome data were mapped against these gene family catologs using diamond (v2.0.4) in blastx mode retaining the best hit per read with at least 50% similarity and default scoring filter options. Mapping counts normalized by gene length were then used to calculate TPM values per gene family. For downstream analyses, rare and low abundant gene families with <= 1 TPM on average and not exceeding 0.1 TPM in at least 20% of the samples were excluded. We used EggNOG to call for functional gene annotations, including KEGG KOs, PFAMs, EC numbers, gene names (“Preferred name”) and CAZy clusters. Abundance profiles of these functional categories were calculated by summing TPM values of genes sharing similar annotations.

### KEGG Module enrichment analysis and cobalamin analysis

The module enrichment analysis was performed from the entire set of KEGG KOs (n = 6,990). The Spearman rank correlation coefficient between KO abundance and PC1 Lifestyle in each country was calculated. Country-level coefficients were averaged using the meta.ave.spear function of vcmeta R package v. 1.4.0 for each KO. Average correlations were then ranked and used as input for the module enrichment analysis, which was performed with the Fast Gene Set Enrichment Analysis (fgsea) R package, version 1.32.4 ^84^, with parameters minSize = 15 and maxSize = 500. The calculation of the p-value depends on randomly generated sets of genes for estimating null-distributions. To assure robust results, the analysis was repeated 100 times by fixing the seed (from one to a hundred) and metrics were only considered for modules with significant adjusted P-values (q-val <0.05, Benjamini-Hochberg correction) in at least 95 out of 100 repetitions. Final metrics (adjusted p-values and normalized enrichment score (NES)) were retrieved from the run with seed = 1.

To identify KEGG KOs primarily driven by lifestyle rather than other host-associated factors, we applied stringent selection criteria. A KO was considered lifestyle-associated only if it met all of the following conditions:

i. Lifestyle or environment showed a statistically significant marginal effect on its variance, as determined by variance partitioning analyses (Fig. 3, Supp. Fig. 3 and Supp. Table 3) (See below the description of the statistical models used for variance partitioning analyses);
ii. The marginal R^2^ for lifestyle exceeded that of both diet and genetics;
iii. The KO abundance profile was significantly correlated with PC1 Lifestyle (Spearman correlation, q-value < 0.05);
iv. The meta-analysis-derived correlation between PC1 Lifestyle and KO abundance at the country level had a standard error that did not span zero;
v. The KO abundance profile was also significantly associated with industrialization status (industrialized vs. non-industrialized, Wilcoxon test, q-value < 0.05).

To analyze the distribution of cobalamin-dependent enzymes in the microbiome, we first retrieved all known bacterial enzymes using cobalamin as a cofactor by querying UniProt with the search term “(cc_cofactor_chebi:“CHEBI:18408”) AND (taxonomy_id:2)” (n = 91,035). We cross-referenced this set with the GMbC protein gene families annotated with a “Preferred name” (gene name) (n = 11,172), yielding 37 proteins. For each of these proteins, we calculated the Spearman correlation between their abundance and PC1 Lifestyle and applied the same variance partitioning approach described below for taxonomic and KEGG features. This allowed us to identify proteins whose abundance was both significantly associated with PC1 Lifestyle and showed a significant marginal variance uniquely attributable to Lifestyle after accounting for all other host factors (Supp. Fig. 3D & Supp. Table 3).

### Detection and profiling of eukaryotic organisms in shotgun metagenomes

We used EukDetect v1.1 ^85^ (--mode runall) and the NBCI taxonomy database to screen shotgun metagenomes for detecting and quantifying eukaryotic taxa.

### Network analyses

Sample matching and screening: We performed three distinct group matching approaches on the metagenomic data to balance both the sample size and the distribution of potential confounders between industrialized and non-industrialized groups. We first performed a matching based on Shannon diversity of TPM-based species-level abundance profiles (Fig. 2) to account for the effect of microbiome diversity on network parameters, using the ‘MatchIt’ R package (v4.5.3), exact matching, and options method=’cem’, ratio=1, caliper=0.1. We also matched groups by age, BMI and sex distributions using the same framework. Finally, we performed matching based on gamma diversity, with the aim of equalizing the number of geographic locations represented in each lifestyle group, as well as the number of samples per location. For this, we considered an iterative process using the ‘entropart’ R package (v1.6-12) and conducted 10,000 random samplings to identify sample sets with the most similar gamma diversity values (gamma diversity, industrialized = 1,214 vs. gamma diversity, non-industrialized = 1,304). Samples from four external cohorts were added to the GMbC shotgun metagenomes to increase the sample size of the industrialized group during sample matching procedures (which inherently reduce sample size in each lifestyle group). We added samples from China (n = 114) ^86^, Germany (n = 60) ^87^, Denmark (n = 50) ^88^ and USA (n = 84, BIO-ML cohort) ^72^. The abundance profiles of these published metagenomes were generated using the same pipeline as for GMbC metagenomes (see above). A full list of samples included in network analyses is provided in Supp. Table 4.

Network construction: We first used FastSpar (v1.0.0) ^40^ to calculate co-abundance microbial correlations and control for sparsity and compositionality of microbiome data. We conserved correlations with FDR < 0.05 and absolute correlation coefficients > 0.3. We then used the ggClusterNet package (v0.1.0) in R to build separate microbial co-abundance networks for industrialized and non-industrialized groups, or for each locality (n >= 30 individuals). To enable comparative analysis of network topology between groups, we calculated the node-level degree centrality in each network using the igraph package (v1.5.1). For standardized comparisons, node degree was normalized by dividing the degree of each node by the maximum possible degree in its respective network, defined as the total number of nodes minus one (N − 1). This normalization accounts for differences in network size between groups, enabling direct comparison of node connectivity patterns across networks.

Natural connectivity: We quantified the structural robustness of microbial interaction networks using a natural connectivity analysis through progressive node removal. The natural connectivity of each intact network was first computed as a baseline measurement of inherent pathway redundancy. To systematically evaluate network stability, we then simulated increasing ecological disturbances by randomly removing nodes in incremental steps. We calculated the natural connectivity at each step, generating a stability profile that captures the resilience of the network to structural perturbations.

Network modules and lineage reconstruction: We considered a layered and ordered graph approach to identify modules and module lineages, with locality-based networks ordered by average position of localities along PC1 Lifestyle. For each locality, we computed the average PC1 Lifestyle position and ordered localities increasingly along this component. All downstream “adjacent localities” refer to consecutive localities in this ordering. Modules were detected on each locality graph using the Leiden algorithm (cluster_leiden in the ‘igraph’ package) with resolution = 0.25 and |r|^3^ as edge weights. Edge sign is ignored during clustering, but preserved for downstream summaries/plots. Every taxon is assigned a module label in its locality. To construct module lineages, we matched modules from each locality to the next along PC1 Lifestyle using the following criteria: 1) modules with >= 4 taxa are eligible for matching and 2) we used the Szymkiewicz–Simpson overlap coefficient to measure taxonomic composition similarity between modules, using similarity >= 0.1 to conserve an eligible pair. We then did a one-to-one assignment of modules. For each adjacent pair of localities, we built the rectangular similarity matrix between eligible source and target modules and solved a linear sum assignment using the cost = 1 - similarity matrix using the Hungarian algorithm (function solve_LSAP from the ‘clue’ R package). For source modules that remain unmatched, we allowed a single-gap lookahead of localities, and tried to match modules to the next adjacent locality with the same similarity threshold and one-to-one assignment. This approach is intended to accommodate for unsampled data and compositional variability across localities. Eligible modules that are not matched at a given locality are allowed to start a new lineage at the next round. We then calculated correlations between lineages and PC1 Lifestyle, correlating module size (taxon count) per locality with PC1 Lifestyle, using Spearman correlation and a standardized slope. Lineages with rho >= 0.30, slope >= 0.05 and p-val <= 0.05 were considered positively correlated to PC1 Lifestyle and ‘emerging’; lineages with rho <= −0.30, slope <= −0.05 and and p-val <= 0.05 were considered disappearing along PC1 Lifestyle. Others were labeled ‘ambiguous’ (Supp. Table 4).

### Differential abundance analyses

Differential abundance analysis (DAA) was performed to identify individual bacterial taxa associated with fecal biomarkers (IgA, IgM, chromogranin, calprotectin). We employed two DAA methods, to validate consistency. We used a standard approach based on multivariate linear regressions of CLR-transformed microbial abundance data. We also used BIRDMAn (v. 0.1.0), a framework for Bayesian modeling that is robust to the high sparsity and compositionality of microbiome data ^89^. Two models were run with each method: one with the baseline covariates (age, sex, BMI, latitude and longitude) and the biomarker of interest, and a second model with the 10 first Lifestyle, Diet and Genetic PCs on top of baseline covariates and biomarker. With BIRDMAn, negative binomial models were fitted. Microbial feature log-fold change was estimated from the resulting posterior distributions, filtered to credible windows excluding zero, and sorted by association with each fecal biomarker.

### Sorting of IgA-coated bacteria and IgA-Sequencing

#### Sample Preparation, Blocking, and IgA Staining

30 mg of each stool sample, homogenized in glycerol, were thawed on ice and transferred to a well of a 2 ml deep-well plate. All stool samples were then washed three times with 500 µL of cold PBS/1% BSA buffer and centrifuged at 4,000 g for 7 minutes at 4°C. Each pellet was then resuspended in 50 µL PBS/1% BSA and 15 µL of mouse serum, and placed on ice for 20 minutes. The total volume of each well was then divided into three separate wells of a deep-well 96-well plate and treated as individual samples for the remainder of the protocol. A 1:40 dilution of anti-human IgA antibody was added to all wells and incubated for 30 minutes on ice. Samples were washed three times with 500 µL of cold PBS/1% BSA buffer and centrifuged at 4,000 g for 7 minutes at 4°C.

#### Bead Binding and Magnetic Separation

We followed a previously-established protocol for magnetic bead-based IgA-Sorting and Sequencing ^50^. Each pellet was resuspended in 100 µL PBS/BSA and stained with 25 µL anti-PE Miltenyi microbeads, then incubated on ice for 30 minutes. The reaction was transferred to a clear polypropylene 96-well plate fitted with a clean plate sheath and a plate magnet on top for 20 minutes to allow magnetic separation. The plate sheath, magnet, and bound samples were then transferred to a wash plate containing 150 µL PBS/1% BSA. The same process was repeated from the wash plate to a final elution plate containing 150 µL PBS. This process was repeated for a total of three 20-minutes magnetic enrichments from the initial plate and three 20-minutes magnetic enrichments from the wash plate.

#### DNA extraction

DNA was then extracted from each of the three triplicates for each sample using the DNeasy UltraClean Microbial kit (Qiagen) immediately after the final enrichment. The samples from the elution plate were labeled as ‘IgA-positive’ fractions, and 15 µL aliquots from each triplicate in the first polypropylene 96-well plate were labeled as ‘IgA-negative’ fractions. DNA was also extracted from the original stool samples of the 300 participants used for IgA sorting. In total, 2,100 samples were processed for 16S sequencing (300 stools + triplicates of the two IgA sorting fractions of these 300 samples).

#### 16S library preparation & sequencing

16S rRNA gene libraries targeting the V4 region of the 16S rRNA gene were prepared for the 2,100 samples by first normalizing template concentrations and determining optimal cycle number via qPCR. Two 25 µL reactions for each sample were amplified with 0.5 units of Phusion with 1X High Fidelity buffer, 200μM of each dNTP, 0.3μM of 515F (5′-AATGATACGGCGACCACCGAGATCTACACTATGGTAATTGTGTGCCAGCMGCCGCGGTAA-3′) and 806rcbc0 (5′-CAAGCAGAAGACGGCATACGAGATTCCCTTGTCTCCAGTCAGTCAGCCGGACTACHVGGGTWTCTAAT-3′). 0.25 µL 100x SYBR was added to each reaction, and samples were quantified using the formula 1.75(ΔCt). To ensure minimal overamplification, each sample was diluted to the lowest concentration sample, amplifying with this sample optimal cycle number for the library construction PCR. Four 25-µL reactions were prepared per sample with master mix conditions listed above, without SYBR. Each sample was given a unique reverse barcode primer from the Golay primer set65. Replicates were then pooled and cleaned via Agencourt AMPure XP-PCR purification system. Purified libraries were diluted 1:100 and quantified again via qPCR (two 25-µL reactions, 2× iQ SYBR SUPERMix (Bio-Rad, ref no. 1708880) with Read 1 (5′-TATGGTAATTGTGTGYCAGCMGCCGCGGTAA-3′), Read 2 (5′-AGTCAGTCAGCCGGACTACNVGGGTWTCTAAT-3′)). Undiluted samples were normalized by way of pooling using the formula mentioned above. Pools were quantified by Qubit (Life Technologies, Inc.) and normalized into a final pool by Qubit concentration and number of samples. Final pools were sequenced on an Illumina MiSeq 300 using custom index 5′-ATTAGAWACCCBDGTAGTCCGGCTGACTGACT-3′ and custom Read 1 and Read 2, mentioned above.

### IgA-Sequencing data analysis

All demultiplexed paired-end 16S read data (n = 2,100 samples) were processed in QIIME2, version qiime2-2019.1. QC and ASV abundance profiling was performed with DADA2 ^90^. Operational Taxonomic Units (OTUs) were reconstructed at 99% similarity. Taxonomic calling was performed using the Greengenes2 trainset (v2022.5, trained with QIIME 2020.8 (sklearn 0.23.1)). We rarefied the 16S read count data to 5,000 reads per sample. We conserved samples that had at least two replicates in each IgA sorting fraction and a 16S stool profile (n = 283). For the analysis of 16S IgA sorting fractions, we filtered out OTUs with less than 10% prevalence in the entire cohort, and OTUs with less than 5% prevalence in at least one of the host lifestyle categories (industrialized vs. non-industrialized). We also filtered out OTUs with less than 0.5% abundance on average across all samples, retaining n = 209 OTUs. OTU x sample combinations for which the OTU was not observed in the stool 16S profile were removed from the analysis. We used the Palm index ^46^ to calculate the IgA-coating index (ICI) of each OTU in each sample. We calculated the average relative abundance across IgA sorting replicates and then calculated the Palm ICI as the ratio between the relative abundance in the positive IgA sorting fraction over the abundance in the negative fraction. We considered an OTU to be coated if ICI > 10. For comparative analyses of IgA coating levels and prevalence between host lifestyle categories (Fig. 6A & B), we controlled for differences in host sample size through subsampling of hosts, and considered OTUs that had ICI data in at least 5 individuals in each category (n = 172).

### Human DNA extraction and genotyping

DNA extraction of human DNA from saliva samples was performed on an automated Chemagic System provided by PerkinElmer. We used the CMG-1091 Nucleic Acid Purification kit from Revvity, with minor adjustments. All samples were genotyped with the Infinium Global Screening Array from Illumina, using 1 µg of DNA. DNA extraction and genotyping was performed at the Genomics platform of the Broad Institute. Genetic variants were called using GRCh37/hg19 as a reference.

### Quality control of GSA data, reconstruction of admixture clusters, genetic PCs and processing of the 1000 human genomes data

The genotyping data was quality controlled with the BIGwas pipeline ^91^. During this step, the GMbC genotypes (n=912) were combined with genotyping data of European population references (ITM (n=3,222) and FOCUS (n=1,004) cohorts ^92^), and with data from two IBD cohorts (EZE & IBD, n=369) of European ancestry ^63^. In short, the pipeline performs detection of the genotyping chip used, renaming and harmonization of variant rs IDs based on the BIGwas annotation database, and filtering out of SNPs deviating from the Hardy-Weinberg equilibrium (false discovery rate (FDR) threshold of 10^-4) or that presented high level of missingness (≥ 0.02 across all batches and ≥0.1 across all batches). Due to the ancestral heterogeneity of the dataset, the SampleQC step of BIGwas was skipped with parameter --skip_sampleqc=1 to avoid removing samples of non-European ancestry. Only variants of the autosomes were used for further analysis. QCed data was imputed with EagleImp-Web ^93^ with the 1000 Genomes Phase 3 reference. The imputed dataset contained 70,314,085 variants with an imputation quality score r² of at least 0.1. The GMbC genotyping data was then combined with data from 1000 Genomes Phase 3 reference. This resulted in a plink fileset, which in the flashpca function of the flashpcaR package for R for PCA reconstruction.

We used ADMIXTURE ^94^ to reconstruct the admixture groups of the GMbC cohort. For this, QCed genotyping data were filtered to include variants with minor allele frequency > 1%. As ADMIXTURE requires unlinked variants, genotype data were LD-pruned using plink and the --- indep-pairwise command (100 kb windows, step size of 50 variants), retaining variants with pairwise r^2^ < 0.05. The resulting 52,060 variants were used to estimate the optimal number of admixture clusters using ADMIXTURE, testing a range of models with K=2 to K=50 clusters, and performing cross-validation. The lowest cross-validation error was observed at K = 17.

### Polygenic risk scores and comparison of allele frequencies

To examine the genetic susceptibility of individuals in the GMbC cohort for IBD, we calculated Polygenic Risk Scores (PRS) and compared these with the PRS of a European IBD cohort, which contains 369 UC patients and 4,281 population controls (1,004 from the FOCUS {and 3,277 from the ITM (Institute for transfusion medicine, UKSH, Kiel) cohorts). Input data included imputed genotyping data, the first ten principal components from BIGwas, and IBD GWAS summary statistics from European individuals ^63^. The summary statistics are based on 5,956 CD and 6,968 UC cases and 21,770 population controls with European ancestry. PRSs were calculated using PRSice v2.5 ^95^ with default parameters. The genotyping data were clumped (250kb regions, clump-p = 1.0, clump-r² = 0.1), and only overlapping variants between the clumped genotyping data and the summary statistics were used for further analysis. To assess the predictive strength of the PRS, different variant sets were tested across variable genome-wide significance thresholds. All GMbC individuals were annotated with a missing phenotype, so that PRSice excludes them from the best model calculation. The optimal model included 13,896 SNPs, yielding a PRS R² of 0.0804.

### Calculation of delta risk allele frequencies for IBD

We used PLINK 1.9 ^96^ on the GMbC data to calculate the allele frequencies of the 232 lead SNPs of the IBD meta-GWAS study ^63^ for each GMbC admixture cluster, and we compared them with the reported frequencies in the IBD meta-GWAS study. The two variants rs5743293 and rs564349 were excluded, as rs5743293 is an indel variant and rs564349 is not present in the 1000 Genomes Phase 3 imputation reference panel. To assess differences, the European frequencies from the IBD meta-GWAS study were subtracted from the observed frequency in each admixture cluster for each variant.

### Measurement of fecal biomarkers

Stool levels of IgA, IgM, chromogranin A (CgA), and calprotectin were measured from samples preserved in glycerol and stored at −80 °C immediately after collection. Measurements were carried out using the following commercial ELISA kits: Secretory IgA ELISA Kit (Immundiagnostik IDK sIgA K8870), Human IgM ELISA Kit (Invitrogen BMS2098), Human Chromogranin AEIA ELISA Kit (Yanaihara Institute Inc. YK070), and Human fecal Calprotectin fCAL ELISA Kit (Bühlmann EK-CAL2). All measurements were done in duplicates. Protein extraction was performed uniformly across all randomly distributed samples. The same extraction supernatant was used for all measurements. All assays were performed according to the manufacturers’ protocols, with adjustments to dilution ratios when necessary.

### Statistical models for variance partitioning

We performed variance partitioning analyses to quantify the individual contributions of lifestyle, diet, and genetics — as well as their non-additive joint effects — to variation in microbiome features and fecal biomarker profiles. We used the first 10 principal components (PCs) from dimensionality reduction of lifestyle, diet, and host genotype data to represent the respective effects of each variable. In the following, the terms Lifestyle, Diet, and Genetics refer to these sets of 10 PCs. We also considered the predictor “Environment”, composed of Lifestyle and Diet.

Variance partitioning was performed with the following response variables: alpha diversity of taxa (Faith’s PD and Shannon) and of KEGG KOs (richness index) (Fig. 2); beta diversity of taxa (first 5 PCs of unweighted UniFrac and Aitchison distances) and of KEGG KOs (first 5 PCs of Canberra distances) (Fig. 2); individual taxa and KEGG KO profiles (Fig. 3); fecal biomarkers (Fig. 5). Analyses were performed on CLR-transformed taxonomic profiles, and on log-transformed KEGG KO abundances.

We defined null and full multivariate linear models, and considered a nested model approach to quantify and test for the significance of marginal and maximum effects of predictors after accounting for baseline host covariates, including Age, Sex, BMI and Geography (latitude and longitude). The following models were defined, and explained variance (R^2^) calculated:

Null model (R^2^_0_), considering baseline host covariates:

Response ∼ Age + Sex + BMI + Latitude + Longitude

Full model (R^2^_Full_), containing the full set of host predictors (baseline covariates + Lifestyle, Diet and Genetics):

Response ∼ Age + Sex + BMI+ Latitude + Longitude + Lifestyle PC1-10 + Diet PC1-10 + Genetics PC1-10

Reduced models compared to the Full model, leaving out one of the non-baseline predictor:

- Without Lifestyle (R^2^_L-_): Response ∼ Age + Sex + BMI + Latitude + Longitude + Diet PC1-10 + Genetics PC1-10
- Without Diet (R^2^_D-_): Response ∼ Age + Sex + BMI + Latitude + Longitude + Lifestyle PC1-10 + Genetics PC1- 10
- Without Genetics (R^2^_G-_): Response ∼ Age + Sex + BMI + Latitude + Longitude + Lifestyle PC1-10 + Diet PC1-10
- Without Lifestyle and Diet (Environment) (R^2^_E-_): Response ∼ Age + Sex + BMI + Latitude + Longitude + Genetics PC1-10

One-predictor extensions of the null model:

- With Lifestyle (R^2^_L+_): Response ∼ Age + Sex + BMI + Latitude + Longitude + Lifestyle PC1-10
- With Diet (R^2^_D+_): Response ∼ Age + Sex + BMI + Latitude + Longitude + Diet PC1-10
- With Genetics (R^2^_G+_): Response ∼ Age + Sex + BMI + Latitude + Longitude + Genetics PC1-10

Calculating the <u>maximum</u> explained variance of all predictors combined, beyond the null model:

R^2^_Max_ = R^2^_Full_ - R^2^_0_

Calculating the <u>maximum</u> explained variance of each predictor:

R^2^_Lmax_ = R^2^_L+_ - R^2^_0_

R^2^_Dmax_ = R^2^_D+_ - R^2^_0_

R^2^_Gmax_ = R^2^_G+_ - R^2^_0_

Calculating the <u>marginal</u> explained variance of each predictor:

R^2^_Lmarg_ = R^2^_Full_ - R^2^_L-_

R^2^_Dmarg_ = R^2^_Full_ - R^2^_D-_

R^2^_Gmarg_ = R^2^_Full_ - R^2^_G-_

Calculating the <u>marginal</u> explained variance of the predictor “Environment”, composed of Lifestyle and Diet:

R^2^_Emarg_ = R^2^_Full_ - R^2^_E-_

Calculating the variance of the non-additive (NA) effect of Diet and Lifestyle:

R^2^_NAmarg_ = R^2^_Emarg_ - R^2^_Lmarg_ - R^2^_Dmarg_

Calculating the variance of the entangled effect of Lifestyle, Diet and Genetics, which is the remainder of the variance not explained by the marginal effects of Genetics and Environment (Diet and Lifestyle, including their non-additive effects):

R^2^_C_ = R^2^_Max_ - R^2^_Gmarg_ - R^2^_Emarg_

### Calculation of the Gut Microbiota Health Index (GMHI)

The Gut Microbiota Health Index (GMHI) is a quantitative metric used to evaluate the health status of the human gut microbiome based on the relative abundance of a curated set of health-associated and disease-associated microbial species ^64^. A positive GMHI indicates enrichment in health-associated taxa, while a negative value reflects a microbiome composition skewed towards disease-associated taxa. A GMHI near zero suggests either a balanced representation of both groups or a lack of detectable marker species. We used the same algorithm and set of marker taxa as described in the original publication to calculate GMHI values ^64^ (https://github.com/jaeyunsung/GMHI_2020). The algorithm uses MetaPhlAn-derived taxonomic profiles. We also calculated MetaPhlAn profiles on metagenomes from the HMP2 IBD cohort (samples from 25 Ulcerative colitis and 39 Crohn’s disease patients) ^97^.

## Data availability

GMbC shotgun metagenomic, IgA-Seq and human genotyping data will be made available online on the dbGaP server (Study ID: 38715; Accession: phs002235.v1.p1; Accession: phs002205.v1.p1) upon publication of the article.

## Code availability

Scripts and command lines used to process the data will be made available at https://github.com/MMmicrobiome-Lab upon publication of the article.

## List of Supplementary materials

## Supplementary Figures

- Supp. Figure 1 – Complementary metadata description of shotgun metagenomes, admixture clusters and dietary drivers of Diet PCs
- Supp. Figure 2 – Impact of host factors and geographic scales on gut microbiome alpha and beta diversity
- Supp. Figure 3 – Associations and variance partitioning of individual microbial taxa and functions with host factors.
- Supp. Figure 4 – Effect of industrialization on co-abundance network characteristics and on the distribution of co-abundance modules
- Supp. Figure 5 – Host and microbial drivers of immune response, gut stress and inflammation biomarkers among GMbC participants.
- Supp. Figure 6 – Conservation and variability of IgA-bacteria binding based on host industrialization status
- Supp. Figure 7 – Low portability of an IBD polygenic risk score to populations of non-European descent.

## Supplementary Tables

- Supp. Table 1 – Participant metadata, admixture clusters and omics statistics
- Supp. Table 2 – Associations and variance partitioning of alpha and beta diversity metrics with host factors
- Supp. Table 3 – Associations and variance partitioning of individual microbial taxa and functions with host factors
- Supp. Table 4 – Co-abundance networks, cohort matching and reconstruction of lineages of co-abundance modules
- Supp. Table 5 – Microbiome-biomarker associations and variance partitioning
- Supp. Table 6 – IgA-Seq binding data and associations with host industrialization
- Supp. Table 7 – Microbiome- and host genome-based predictors of diseases

## Supplementary Information

- Weak effects of recent antibiotic/antiparasite consumption on alpha and beta microbiome variation.
- Definition and assignment of subsistence strategies

### Acknowledgments

This work was supported by grants from the Center for Microbiome Informatics and Therapeutics at MIT and the Rasmussen Family Foundation.

M.P, M.R and M.G. received support from the Deutsche Forschungsgemeinschaft (DFG - German Science Foundation) within the Collaborative Research Center (CRC) 1182 on “The Origin and Function of Metaorganisms” (Project-ID 261376515 – SFB 1182, project C5.1 to M.G., project C5.2 to M.P., project C5.3 to M.R.).

The study received infrastructure support from the DFG (German Science Foundation) within the Cluster of Excellence 2167 “PrecisionMedicine in Chronic Inflammation (PMI)” (EXC 2167-390884018) and the DFG research unit “miTarget” (project number 426660215; INF (EL 831/5-2)).

M.G. received funding from the European Research Council (ERC) under the European Union’s Horizon 2020 research and innovation programme (CoG VESICULOME, Grant agreement No. 101126254).

M.G., M.P. and A.F. received funding from the DFG – Project number 426660215 with the RU 5042 miTarget on “The microbiome as a therapeutic target in inflammatory bowel disease”, subprojects TP01 “Targeting intestinal yeasts and pathogenic yeast-responsive CD4+ T cells in Crohn’s disease” and TP02 “Ecology and function of synthetic bacterial communities for the understanding and modulation of IBD-associated microbiomes”.

The project received funding from the European Union under the Horizon Europe grant agreement No. 101095470 (project miGut-Health). Views and opinions expressed are however those of the author(s) only and do not necessarily reflect those of the European Union nor European Health and Digital Executive Agency (HaDEA). Neither the European Union nor HaDEA can be held responsible for them.

A.P.S. is funded by a grant of KiTE-Kiel Training for Excellence. The project KiTE – Kiel Training for Excellence (KiTE) has received funding from the European Union’s Horizon Europe research and innovation programme under the Marie Skłodowska-Curie grant agreement No 101081480.

L.P. is supported by the University of California San Diego Medical Scientist Training Program (NIH/NIGMS T32GM007198).

A.S. was supported by a Marie Sklodowska-Curie Actionsfellowship (H2020-MSCA-IF-2016-780860, H2020-MSCA-IF-2020-101032316).

A.Z acknowledges support from the University of Strasbourg’s Initiative of Excellence.

L.S. was supported by an Agence nationale de la recherche (ANR) grant (MICROREGAL, ANR-15-CE02-0003).

K.A.V. and L.R.R. were supported by the Mayo Clinic Center for Clinical and Translational Science (TL1 TR002380 and UL1 TR002377, respectively).

B. J. S. acknowledges funding from the Canada Research Chairs Program.

R. J. X. acknowledges funding from the National Institutes of Health (NIH) (Project Center for the Study of Inflammatory Bowel Disease at Massachusetts General Hospital - DK043351)

We thank Tamara Mason and the team at the Walkup Sequencing platform, as well as the Infectious Disease and Microbiome Program’s Microbial Omics Core at the Broad Institute of MIT and Harvard for their support on sequencing efforts.

We thank Kiel Life Science, the Kiel Microbiome Center and the NFDI4Microbiota project (Flex Fund grant, DFG project no. 28/1 AOBJ: 700895 Bio4ALL, NFDI4Microbiota) as part of the German National Research Data Infrastructure (NFDI) for supporting outreach and capacity building activities of the GMbC.

We are grateful to all communities and relevant stakeholders for authorizing this work, and to the Polar Continental Shelf Program (Natural Resources Canada) for logistical support and lab space.

## Author contributions

Designed this study: M.G, M.P, M.R, E.J.A

Field administrative work & collection of data and samples: M.P, M.G, L.T.T.N, A.Z, A.A, M.Y.A, S.O.A, Y.A.A, A.S.B, L.S.C, C.C, A.DE, M.D, A.DU, A.FE, A.FR, S.G, C.G, J.H, F.I, D.I, V.J, P.K, S.L, E.L, J.L, Y.A.L.L, A.M, V.M, R.S.M, I.E.M, Y.A.N, D.N, M.N, C.O, T.M.P, A.P, L.R, L.RU, J.R, L.S, B.J.S, S.S, A.S, K.V, T.V, R.V, K.M

Performed processing of biospecimens and molecular work: M.P, M.G, L.T.T.N, C.S, J.C

Biosample and Data curation: M.P, M.G, M.R, L.M-S, H.J

Data analysis: M.G, M.P, M.R, A.P.S, Y.M, L.M-S, E.M.W, L.P, D.P, D.M

Supervision: M.G, M.P, E.J.A

Funding acquisition: M.P, M.G, E.J.A, R.J.X, M.D

Writing, original draft: M.G, M.P, M.R

Writing, review & editing: all authors

---

**Supplementary Figure 1 – Complementary metadata description of shotgun metagenomes, admixture clusters and dietary drivers of Diet PCs**

A. Distribution of mapping rates against SGB reference genomes (top) and 50% protein gene families (bottom). Mapping rates are grouped by country of origin. Intervals show standard deviations of per-country mapping rates. The Salmon and Diamond tools were used for mapping short metagenomic reads against reference databases.

B. Posterior probabilities of admixture clusters across GMbC participants. Participants are grouped by country and self-declared ethnic groups. Admixture cluster reconstructions were performed with ADMIXTURE.

C. Spearman correlations between Diet PCs (rows) and Lifestyle PCs (columns, left panel) or individual dietary items (columns, right panel). Correlation coefficients are colored-coded along a gradient from red to blue. Significant correlations (adjusted p-values) are shown as plain points.

**Supplementary Figure 2 – Impact of host factors and geographic scales on gut microbiome alpha and beta diversity**

A. Correlations between Faith’s phylogenetic diversity metric and PC1 Lifestyle, per country. Linear regression lines for each country are shown. Lines are colored by country (color legend below). Statistics of spearman correlations are shown in Supp. Table 2. All regressions have negative correlation coefficients, to the exception of Finland. Correlations are significant for Cameroon, Ghana, Malaysia, Rwanda, Senegal and Thailand.

B. Population density is strongly associated with shifts in microbiome compositions. The panel depicts a PCoA of unweighted UniFrac compositional dissimilarities at the species level. Individual participants are shown in the background and are colored in grey. Average positions of sampled localities are shown. Localities are colored by population density (log scale).

C. Distribution of inter-individual microbiome compositional dissimilarity (unweighted UniFrac) based on geographical distance (x axis, distances in km). Geographic distances were calculated with the Haversine distance metric, which calculates the shortest distance between two points over the surface of a sphere. The blue line represents the lineage regression. Spearman correlation, ρ = -0.04, p = 1.8e-138

D. PCoA of KEGG KO profiles. KEGG KO dissimilarities were calculated with the Canberra distance metric. Participants are colored by country as in A. PERMANOVA test with PC1 Lifestyle: R2 = 0.078, p-val < 0.001.

E. PCoA of 50% protein family profiles. Dissimilarities were calculated with the Bray-Curtis distance metric. Participants are colored by country as in A. PERMANOVA test with PC1 Lifestyle: R2 = 0.08, p-val < 0.001

**Supplementary Figure 3 – Associations and variance partitioning of individual microbial taxa and functions with host factors.**

A. Full results of the meta-analysis of country-level associations between gut bacterial species abundance and urbanism status (urban vs. rural) shown in Fig. 3A. The “META-ANALYSIS” column shows statistical results of the cross-country meta-analysis performed using an inverse-variance weighted fixed-effects approach. Because Canada included only rural participants in our cohort, it was grouped with the USA under the label ‘North America’. Species are shown in rows.

B. Effect of host covariates on the association between bacterial genera abundance and industrialization (PC1 Lifestyle) (x axis: genera; y axis: effect size). Two models were compared – a first model with Lifestyle PCs alone as predictors, and a second model with host covariates (baseline covariates and Diet and Genetic PCs). Significant associations are shown as plain symbols.

C. Variance partitioning of KEGG KO abundance to calculate marginal and maximum variance of baseline, Lifestyle, Diet and Genetic factors, along with Non-additive Effects. The same statistical framework and figure design as in Fig. 3C were used. “Environment” is defined as the overall contribution of Lifestyle and Diet (see Methods). Features are grouped by majority factors that explain >50% of variance. Top 50 hits per majority factor are shown. Full results are shown in Supp. Table 3.

D. Correlation of the abundance of cobalamin-dependent enzymes in the microbiome and host lifestyle. Left: correlation with PC1 Lifestyle. Positive correlations (Spearman correlation) are shown in purple and indicate higher abundance in more industrialized host populations. Negative correlations are shown in green and indicate higher abundance in more non-industrialized populations. Right: Marginal variance (R2) of Lifestyle explaining the abundance distribution of each enzyme. The variance partitioning approach used is the same as the one used on taxonomic and KEGG KO profiles (see Methods). Plain bars indicate significant marginal variance.

**Supplementary Figure 4 – Effect of industrialization on co-abundance network characteristics and on the distribution of co-abundance modules**

A. Age distribution between industrialized and non-industrialized groups after matching cohorts for age.

B. Same as in A, for BMI.

C. Same as in A, for Shannon alpha diversity of microbial taxa.

D. Same as A, for host sex.

E. Same as in A, for number of localities and per-locality number of participants.

F. Increased node degree among co-abundance networks of more industrialized populations when considering lifestyle groups with participants unmatched for potential confounders. Wilcoxon test.

G. Increased node degree among co-abundance networks of more industrialized populations after matching cohorts for BMI, age and sex. Wilcoxon test. Median node degree, industrialized = 0.0119; non-industrialized = 0.00819.

H. Increased node degree among co-abundance networks of more industrialized populations after matching cohorts for participant distribution across geographic localities. Wilcoxon test. Median node degree, industrialized = 0.0137; non-industrialized = 0.0131.

I. Module lineages with significant positive/negative trends along PC1 Lifestyle. Localities (x-axis) are ordered by mean PC1 Lifestyle; lineages are shown in y-axis (see Supp. Table 4). The heatmap shows the presence and size of the module in a given locality. Size of modules (number of taxa) is represented along a color gradient. Modules were matched across adjacent localities using taxonomic compositional overlap (Szymkiewicz– Simpson), with one-to-one assignment (Hungarian) and a single-gap lookahead to accommodate for module absence. Top lineages are negatively associated with PC1 Lifestyle (markers of less industrialized contexts); bottom lineages are positively associated (markers of more industrialized contexts).

J. Label-swap permutation to test whether the number of positively and negatively associated lineages with PC1 Lifestyle is higher than expected under a null distribution. The permutation randomly re-assigned locality PC1 values, and module detection and lineage reconstructions were calculated for each replicate. Observed counts (dashed lines) are higher than null expectations (p < 0.001 and p = 0.002 for negatively (top panel) and positively-associated lineages (bottom panel), respectively).

**Supplementary Figure 5 – Host and microbial drivers of immune response, gut stress and inflammation biomarkers among GMbC participants.**

A. Correlations among IgA, IgM, Calprotectin and Chromogranin levels, correcting for age, sex, BMI, Latitude, Longitude and Genetic PCs (linear model). Statistical significance is based on Bonferroni-corrected p-values.

B. Elevated fecal IgA among urban communities. Fecal IgA was compared between urban (blue) and rural (red) participants in countries where both groups are represented. Countries are ordered by mean value along PC1 Lifestyle. Wilcoxon test. Significance is as follows: *: p < 0.05; **: p < 0.01; ***: p < 0.001.

C. Higher levels of detected Eukaryotes among non-industrialized participants. The panels show the distribution of detected eukaryotes in metagenomes across major eukaryotic taxonomic groups and host lifestyles.

D. Association between Eukaryote presence/absence profiles in the metagenomes and fecal biomarkers. ‘Euk Pres/Abs’ represents the aggregated presence/absence profiles across all eukaryotic taxa. The marginal variance explained by eukaryotic features was quantified using a nested-model approach, with each biomarker as the response variable in a linear model including age, sex, BMI, latitude, longitude, lifestyle PCs, diet PCs, genetic PCs, and microbiome PCs (unweighted UniFrac) as predictors.

E. Correlation between fecal IgA levels (y axis) and the abundance of IgA peptidases in the metagenome, defined as the sum of TPM counts across all 50% gene families annotated as IgA peptidases. Linear regressions for each lifestyle category are shown (industrialized: beta = 0.6194, p-val = 0.04; non-industrialized: beta = 0.7883, p-val = 1.45e-07).

F. Consistent association between microbiome compositions and fecal IgA across countries. PERMANOVA tests were run between unweighted UniFrac-based microbiome dissimilarities and IgA levels for each country. Significance is as follows: *: p < 0.05; **: p < 0.01; ***: p < 0.001.

A. G. Comparison of fecal Chromogranin levels between urban (blue) and rural (red) participants in countries where both groups are represented. Countries are ordered by mean value along PC1 Lifestyle. Wilcoxon test. Significance is as follows: *: p < 0.05; **: p < 0.01; ***: p < 0.001. Higher levels of Chromogranin among urban communities were detected in Central African Republic, Rwanda, Malaysia, Ghana and Thailand.

B. H. Density distribution of participants with moderate-to-high levels of fecal Calprotectin (> 50ug/g stool) along PCoA1 of microbiome compositions (unweighted UniFrac). See Fig. 2B for the distribution of participants, geographies and host lifestyles along PCoA1. The density distribution suggests an increase in frequency of participants with moderate-to-high Calprotectin levels among urban and industrialized participants.

I. Distribution of participants per locality with Calprotectin levels >= 50ug/g stool and at least one (left panel) or two (right panel) calprotectin-associated pathobionts detected in the metagenome. Pathobionts considered are E. faecalis, E. coli and K. pneumoniae.

**Supplementary Figure 6 – Conservation and variability of IgA-bacteria binding based on host industrialization status.**

A. Correlation of the prevalence of OTU-specific IgA coating (ICI > 10) based on industrialization status. The linear regression, credibility interval and associated statistics are shown.

B. Comparison of the frequency of IgA-coated genera (ICI > 10) by industrialization (left) and urbanism (right) status. Genus-level ICI values were calculated as the summed counts of all OTUs within each genus in the positive and negative IgA fractions.

C. Two example OTUs for which the prevalence of IgA coating (ICI > 10) is elevated in the industrialized group.

**Supplementary Figure 7 – Low portability of an IBD polygenic risk score to populations of non-European descent.**

A. Distribution of the IBD polygenic risk score (PRS) across GMbC admixture clusters with >45 individuals. For comparison, the distributions of IBD PRS from two external European cohorts (one healthy, one with IBD) are also shown (see Methods).

B. Distributions of delta risk allele frequency (%) of IBD risk alleles used to calculate the PRS across admixture clusters, showing different allele frequency spectra in groups of non-European descent. Patterns in A and B are consistent with known limitations in PRS portability across ancestries, reflecting differences in allele risk frequencies (among other factors), rather than true differences in genetic risk.

